# Hypoxia truncates and constitutively activates the key cholesterol synthesis enzyme squalene monooxygenase

**DOI:** 10.1101/2022.08.18.504470

**Authors:** Hudson W. Coates, Ellen M. Olzomer, Ximing Du, Rhonda Farrell, Hongyuan Yang, Frances L. Byrne, Andrew J. Brown

## Abstract

Cholesterol synthesis is both energy- and oxygen-intensive, yet relatively little is known of the regulatory effects of hypoxia on pathway enzymes. We previously showed that the rate-limiting and first oxygen-requiring enzyme of the committed cholesterol synthesis pathway, squalene monooxygenase (SM), can undergo partial proteasomal degradation that renders it constitutively active. Here, we show that hypoxia is the physiological trigger for this truncation, which occurs through a two-part mechanism: (1) increased targeting of SM to the proteasome *via* stabilization of the E3 ubiquitin ligase MARCHF6, and (2) accumulation of the SM substrate, squalene, which impedes the complete degradation of SM and liberates its truncated form. Truncation of SM is also increased in endometrial cancer tissues, where it correlates with levels of hypoxia-inducible factor−1α. These results uncover a feedforward mechanism that enables SM to accommodate fluctuations in substrate levels yet is also a likely contributor to its widely reported oncogenic properties.

## Introduction

Cholesterol is an essential component of mammalian cell membranes, yet its aberrant accumulation is detrimental [1]. Most cellular cholesterol arises from an energetically expensive biosynthetic pathway requiring eleven oxygen molecules and over one hundred ATP equivalents per molecule of product [2]. Furthermore, many intermediates of this pathway are toxic in excess [3]. Coordinated regulation of cholesterol synthesis enzymes is therefore vital to ensure that the pathway is active only when required, and that sufficient substrates and cofactors are available to maintain flux through the full length of the pathway.

Squalene monooxygenase (SM, also known as squalene epoxidase or SQLE, EC:1.14.14.17) catalyzes the rate-limiting conversion of squalene to monooxidosqualene in the committed cholesterol synthesis pathway [4]. This reaction is the first of the pathway to require molecular oxygen, with the introduced epoxide group ultimately forming the signature C3-hydroxyl group of cholesterol. SM can also act a second time on monooxidosqualene to produce dioxidosqualene, the precursor of the potent regulatory oxysterol 24(*S*),25-epoxycholesterol [5]. As a flux-controlling enzyme, SM is subject to metabolic regulation at both the transcriptional level *via* sterol regulatory element-binding proteins [6] and the post-translational level *via* ubiquitination and proteasomal degradation [4]. The latter is mediated by the N-terminal regulatory domain of SM (SM-N100), which senses lipid levels in the endoplasmic reticulum (ER) membrane and accelerates or attenuates SM degradation in response to excess cholesterol or squalene, respectively [7, 8]. These reciprocal feedback and feedforward loops thus fine-tune SM activity according to metabolic demand. SM is typically fully degraded by the proteasome; however, incomplete proteolysis produces a truncated form of SM (trunSM) that lacks a large portion of the lipid-sensing SM-N100 domain but retains the full catalytic domain [9]. This renders trunSM cholesterol-resistant and therefore constitutively active. Although truncation is induced by the SM inhibitor NB-598, human cell lines express similar levels of full-length and truncated SM [9]. This points to the existence of an unknown physiological trigger for truncation.

Clarifying the mechanisms of SM regulation is particularly pertinent given the importance of the enzyme, and cholesterol more generally [10], in oncogenesis. Overexpression of the SM gene *SQLE* is associated with greater invasiveness and lethality in breast [11], prostate [12, 13], and pancreatic cancer [14], amongst others. *SQLE* also shares its genomic locus with the oncogene *MYC*, with which it is frequently co-amplified in cancer [11, 15]. Moreover, myc itself transcriptionally upregulates *SQLE* expression [16], while the tumor suppressor protein p53 downregulates its expression [17]. At the protein level, aberrant SM expression is implicated in colorectal cancer progression [18, 19] and the development of both nonalcoholic steatohepatitis and hepatocellular carcinoma [20, 21]. Given its key role in oxygen-dependent cholesterol synthesis, SM may be particularly critical for cancer cell survival during hypoxia, which is common in the poorly vascularized cores of solid tumors and often associated with poor prognosis [22]. In support of this idea, SM inhibition sensitizes breast and colorectal cancer cells to hypoxia-induced cell death [23]. Although hypoxic cells tend to accumulate cholesterol, there are conflicting reports on changes in biosynthetic flux [24–26]. Furthermore, with the notable exception of the early pathway enzyme 3-hydroxy-3-methylglutaryl-CoA reductase (HMGCR) [27], the effects of hypoxia on individual biosynthetic enzymes are unknown. It is also unclear whether these might be perturbed in a tumor context to favor continued cholesterol synthesis and cell proliferation.

Here, we show that hypoxic conditions induce the truncation of SM in a variety of cell lines through a combination of accelerated proteasomal degradation and inhibition of its complete proteolysis. This occurs due to the accumulation of both MARCHF6, the major E3 ubiquitin ligase for SM, and squalene, which impedes SM degradation through a mechanism involving the SM-N100 regulatory domain. We also show that SM truncation is increased and correlates with the magnitude of hypoxia in endometrial cancer tissues. Taken together, our findings point towards a role for the constitutively active trunSM in adaptations to hypoxic conditions and suggest that it contributes to the oncogenic effects of SM activity.

## Results

### Oxygen availability regulates SM truncation

We previously showed that SM is post-translationally regulated by both its substrate, squalene, and its pathway end-product, cholesterol [4, 8]. The enzyme also undergoes partial proteasomal degradation of its N-terminus to liberate a truncated protein (trunSM) that is cholesterol-resistant and thus constitutively active (Fig. 1A) [9], although the physiological trigger is unknown. As SM is a rate-limiting enzyme of cholesterol synthesis and catalyzes its first oxygen-dependent reaction, we next tested if SM protein levels are affected by oxygen availability. Incubation of HEK293T cells under hypoxic conditions (1% O_2_) stabilized hypoxia-inducible factor-1α (HIF1α; Fig. 1B) and upregulated its target genes *VEGF* and *CA9* (Supplementary Fig. S1A), confirming the induction of a hypoxic response. We also noted a striking increase in SM truncation under these conditions, as indicated by the disappearance of full-length SM and a four-fold accumulation of trunSM (Fig. 1B). Hypoxia-induced truncation of SM increased over time (Fig. 1C) and with the magnitude of oxygen deprivation (Fig. 1D). Notably, trunSM accumulation was greater under the severely hypoxic conditions characteristic of solid tumors (0.5–2% O_2_) than the ‘physoxic’ conditions experienced by normal human tissues *in situ* (3–7.5% O_2_) [28]. This suggested that increased SM truncation is a feature of pathophysiological hypoxia. The net result of hypoxia-induced truncation was a progressive increase in the levels of total SM over time (expressed as the sum of full-length SM and trunSM levels; Supplementary Fig. S1B).

**Figure 1.**
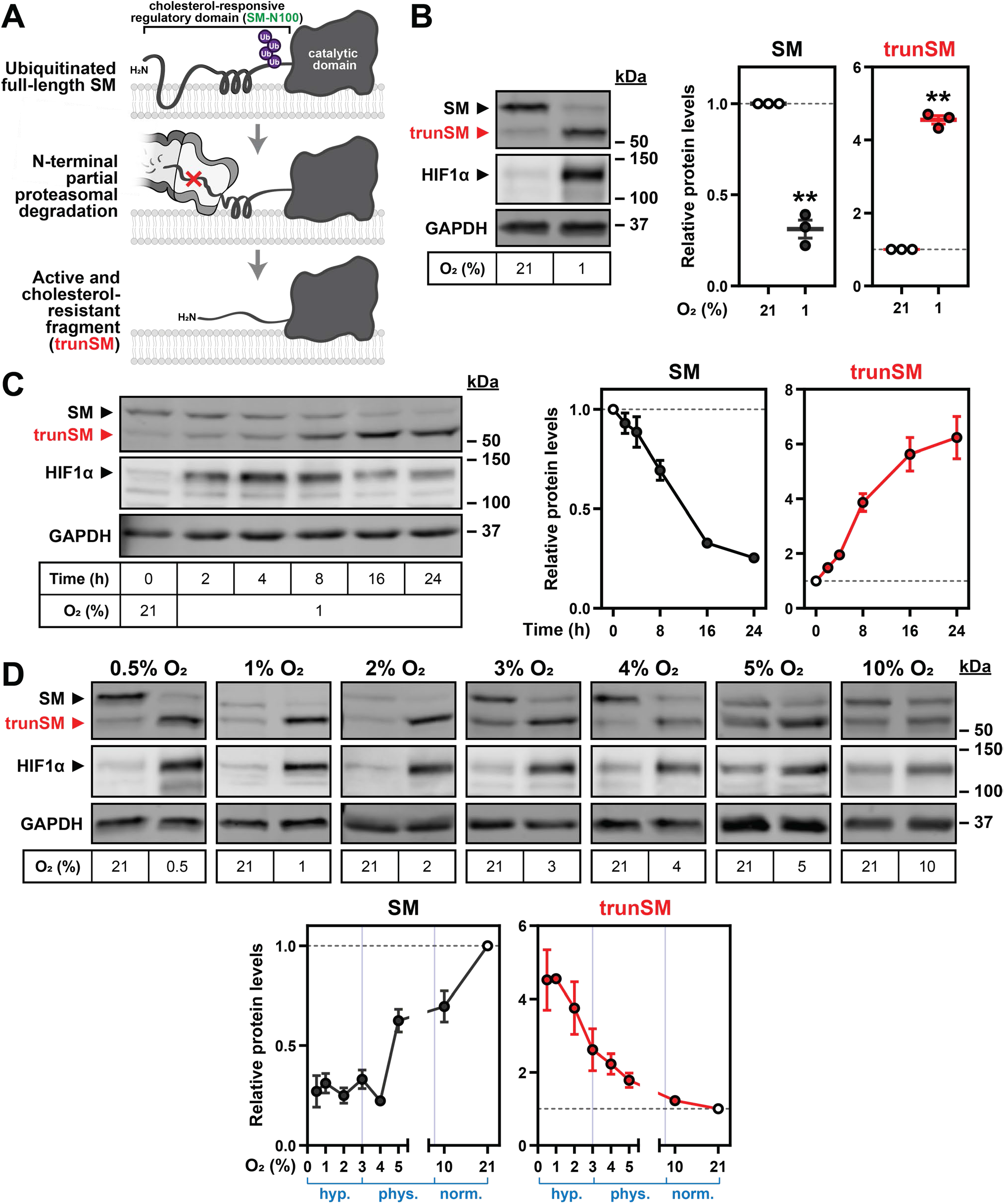
Oxygen availability regulates SM truncation. **(A)** Simplified overview of SM truncation [9]. Full-length SM contains an N-terminal domain responsible for feedback regulation by cholesterol. Ubiquitinated SM is targeted to the proteasome, where proteolysis is prematurely halted within the N-terminal regulatory domain. This liberates a truncated protein (trunSM) that no longer responds to cholesterol and is therefore constitutively active. **(B)** HEK293T cells were incubated under normoxic (21% O_2_) or hypoxic (1% O_2_) conditions for 24 h. **(C)** HEK293T cells were incubated under normoxic or hypoxic conditions for the indicated times. Changes in HIF1α levels over time are consistent with other reports [29, 30]. **(D)** HEK293T cells were incubated under the indicated oxygen concentrations (0.5–21% O_2_) for 24 h. Each set of immunoblots was obtained in a separate experiment. **(B–D)** Immunoblotting was performed for SM and truncated SM (trunSM, red). Graphs depict densitometric quantification of protein levels normalized to the normoxic condition, which was set to 1 (dotted line). In (D), oxygen concentrations considered hypoxic (Hyp.), ‘physoxic’ (Phys.) or normoxic (Norm.) [28] are indicated in blue. Where error bars are not visible, variance is too low to be depicted. Data presented as mean ± SEM from *n* ≥ 3 independent experiments (**, *p* ≤ 0.01; two-tailed one- sample *t*-test vs. hypothetical mean of 1).

We also surveyed SM levels in a panel of cell lines and found that hypoxia-induced accumulation of trunSM was generalizable to all, although full-length SM levels did not decrease in MDA-MB-231 breast cancer cells (Supplementary Fig. S1C). As HIF1α and hypoxia-inducible factor-2α (HIF2α) transcriptionally regulate the cellular response to hypoxia, we next tested if their activity is required for SM truncation. However, knockdown of *HIF1A* and *HIF2A* gene expression in HEK293T cells had no effect on the hypoxia- induced truncation of SM (Supplementary Fig. S1D, E), ruling out the involvement of these transcription factors and their target genes in the phenomenon.

### Hypoxia transcriptionally and post-translationally reduces full-length SM levels

As the truncation of SM results from its partial proteasomal degradation [9], we reasoned that hypoxia promotes truncation through a two-part mechanism: (1) targeting of full-length SM to the proteasome, and (2) inhibition of its complete degradation. To confirm the first step of this mechanism, we investigated the reason for the decline in full-length SM levels during hypoxia. *SQLE* transcripts were downregulated in hypoxic HEK293T cells, as were transcripts encoding the upstream cholesterol and isoprenoid synthesis enzyme HMGCR (Fig. 2A). Downregulation of *SQLE* transcripts was not observed in MDA-MB-231 cells (Supplementary Fig. S2A), accounting for the unchanged full-length SM levels in this cell line. Although the reduction in *SQLE* and *HMGCR* transcripts in HEK293T cells likely reflected a broad transcriptional suppression of cholesterol synthesis during hypoxia, as reported previously [31, 32], the magnitude of *SQLE* downregulation was unlikely to fully explain the large reduction in SM protein levels (Fig. 1B). Moreover, levels of a constitutively expressed SM construct were markedly reduced during extended hypoxic incubations, with no associated changes in mRNA levels (Fig. 2B, Supplementary Fig. S2B). We concluded that hypoxia reduces the levels of full-length SM through both transcriptional downregulation and accelerated post-translational degradation.

**Figure 2.**
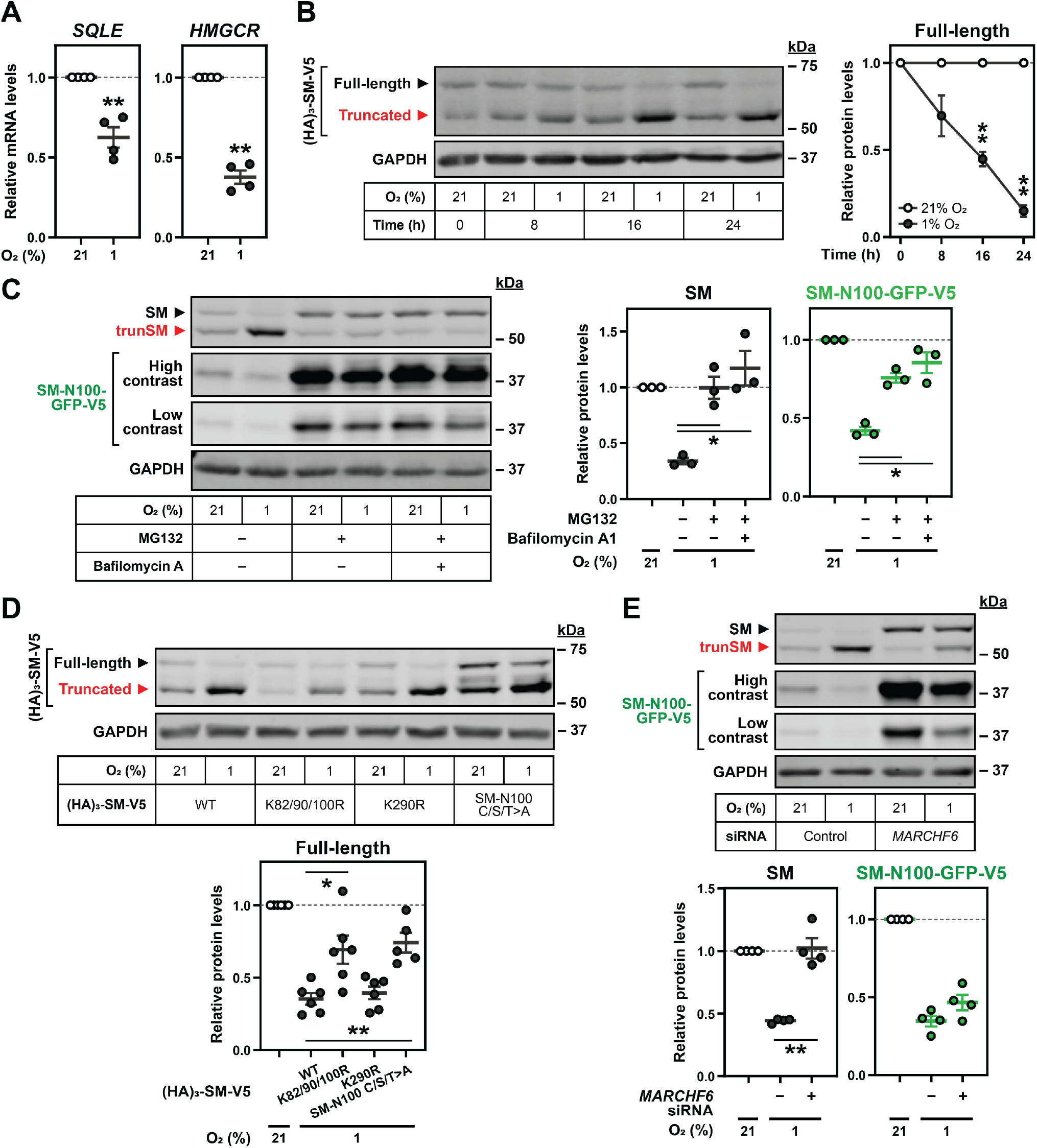
Hypoxia transcriptionally and post-translationally reduces full-length SM levels. **(A)** HEK293T cells were incubated under normoxic or hypoxic conditions for 24 h. Levels of the indicated transcripts were quantified, normalized to the levels of three housekeeping transcripts and adjusted relative to the normoxic condition, which was set to 1 (dotted line). **(B)** HEK293T cells were transfected with (HA)_3_-SM-V5 for 24 h and incubated under normoxic or hypoxic conditions for the indicated times. **(C)** HEK SM-N100-GFP-V5 cells were treated with or without 20 μM MG132 and 20 nM bafilomycin A1 under normoxic or hypoxic conditions for 16 h. **(D)** HEK293T cells were transfected with the indicated constructs for 24 h and incubated under normoxic or hypoxic conditions for 16 h. **(E)** HEK SM-N100-GFP-V5 cells were transfected with control or *MARCHF6* siRNA for 24 h and incubated under normoxic or hypoxic conditions for 16 h. **(B–E)** Graphs depict densitometric quantification of protein levels normalized to the respective normoxic conditions, which were set to 1 (dotted line). **(A–E)** Data presented as mean ± SEM from *n* ≥ 3 independent experiments (*, *p* ≤ 0.05; **, *p* ≤ 0.01; [A, B] two-tailed one-sample *t*-test vs. hypothetical mean of 1; [C–E] two-tailed ratio paired *t*-test vs. vehicle, wild-type [WT] or control siRNA condition).

The basal and metabolically-regulated degradation of SM occurs through the ubiquitin-proteasome system and is mediated by the SM-N100 regulatory domain [7, 8]. Therefore, we tested the effect of hypoxia on HEK293 cells stably overexpressing an SM-N100 fusion protein (SM-N100-GFP-V5). Like full-length SM, levels of SM-N100-GFP-V5 were reduced under hypoxic conditions (Fig. 2C). Proteasomal inhibition using MG132 rescued SM and SM-N100-GFP-V5 from this reduction, confirming that hypoxia-induced degradation occurs *via* the proteasome. We also noted that the hypoxia-induced accumulation of trunSM was ablated by MG132 (Supplementary Fig. S2C), consistent with the protein arising from partial proteasomal proteolysis of SM [9]. Although hypoxia can trigger autophagy [33], this did not play a role in SM degradation as inhibition of lysosomal acidification using bafilomycin A1 had no additive effect with MG132 (Fig. 2C). To identify residues required for hypoxia-induced degradation of SM, we utilized protein constructs with mutations of previously identified ubiquitination sites. Degradation was blunted by the disruption of Lys-82/90/100, a cluster of redundant ubiquitination sites previously found to promote truncation [9], but not by the disruption of Lys-290 [34] (Fig. 2D). Non-canonical cysteine, serine and threonine ubiquitination sites required for the cholesterol-induced degradation of SM [35] also contributed to hypoxia-induced degradation, suggesting that multiple ubiquitin signals are involved.

To investigate the mechanism by which hypoxia promotes SM ubiquitination, we considered the possible role of proline hydroxylation. This oxygen-dependent modification, catalyzed by prolyl hydroxylases, is required for the ubiquitination and degradation of HIF1α under normoxic conditions [36], although there is conflicting evidence for the existence of substrates beyond the HIF proteins [37]. Indeed, treatment with prolyl hydroxylase inhibitors had no effect on SM nor SM-N100 levels despite stabilizing HIF1α (Supplementary Fig. S2D). SM and SM-N100 are targeted for proteasomal degradation by the E3 ubiquitin ligase MARCHF6 [38]; therefore, we next tested if increased MARCHF6 activity could account for the hypoxia-induced degradation of SM. To do so, we depleted *MARCHF6* expression using siRNA, which we have previously shown achieves a 60–70% reduction in transcript levels in this cell line [38]. *MARCHF6* knockdown reversed the hypoxic decline in full-length SM levels (Fig. 2E), supporting its involvement in hypoxia-induced degradation. The relative accumulation of trunSM was also blunted (Supplementary Fig. S2E), consistent with truncation occurring post-ubiquitination by MARCHF6. This effect was not abolished, however, indicating that SM can be truncated even when targeted to the proteasome by other mechanisms. Surprisingly, there was no significant effect of *MARCHF6* knockdown on hypoxia-induced degradation of SM-N100-GFP-V5 (Fig. 2E), suggesting that full-length SM and the isolated SM-N100 domain are degraded through different proteasome-dependent routes under these conditions. As truncation involves the full-length protein [9], we elected to further investigate the MARCHF6-mediated regulation of SM.

### Hypoxia stabilizes the E3 ubiquitin ligase MARCHF6

To study MARCHF6 levels in hypoxic cells, we utilized a previously generated HEK293 cell line stably overexpressing a V5-tagged form of the protein. This construct was used due to the poor performance of endogenous MARCHF6 antibodies [39], and also to eliminate any transcriptional contribution to protein levels. We first examined the response of MARCHF6-V5 to hypoxia and found that it accumulated during prolonged hypoxic incubations, correlating with the maximal decline in full-length SM levels (Fig. 3A). Previously, we showed that cholesterol stabilizes MARCHF6 by interfering with its autoubiquitination and proteasomal degradation [39]. To determine if a similar mechanism accounted for the accumulation of MARCHF6 in hypoxia, we treated MARCHF6-V5 cells with CB-5083, an inhibitor of the ER-associated degradation cofactor valosin-containing protein, or the proteasome inhibitor MG132. Both treatments stabilized MARCHF6, with CB-5083 having a greater effect, as described previously [39]. Importantly, both CB-5083 and MG132 blocked the hypoxia-induced accumulation of MARCHF6 (Fig. 3B), consistent with a mechanism involving reduced proteasomal degradation. We also tested the effect of hypoxia on two MARCHF6 mutants: C9A, an inactive variant lacking a functional RING domain [40], or N890A, a variant that is defective in auto-ubiquitination [41]. Both mutants showed reduced accumulation during hypoxia (Fig. 3C), indicating that the effect on wild-type MARCHF6-V5 is due to impaired auto-ubiquitination.

**Figure 3.**
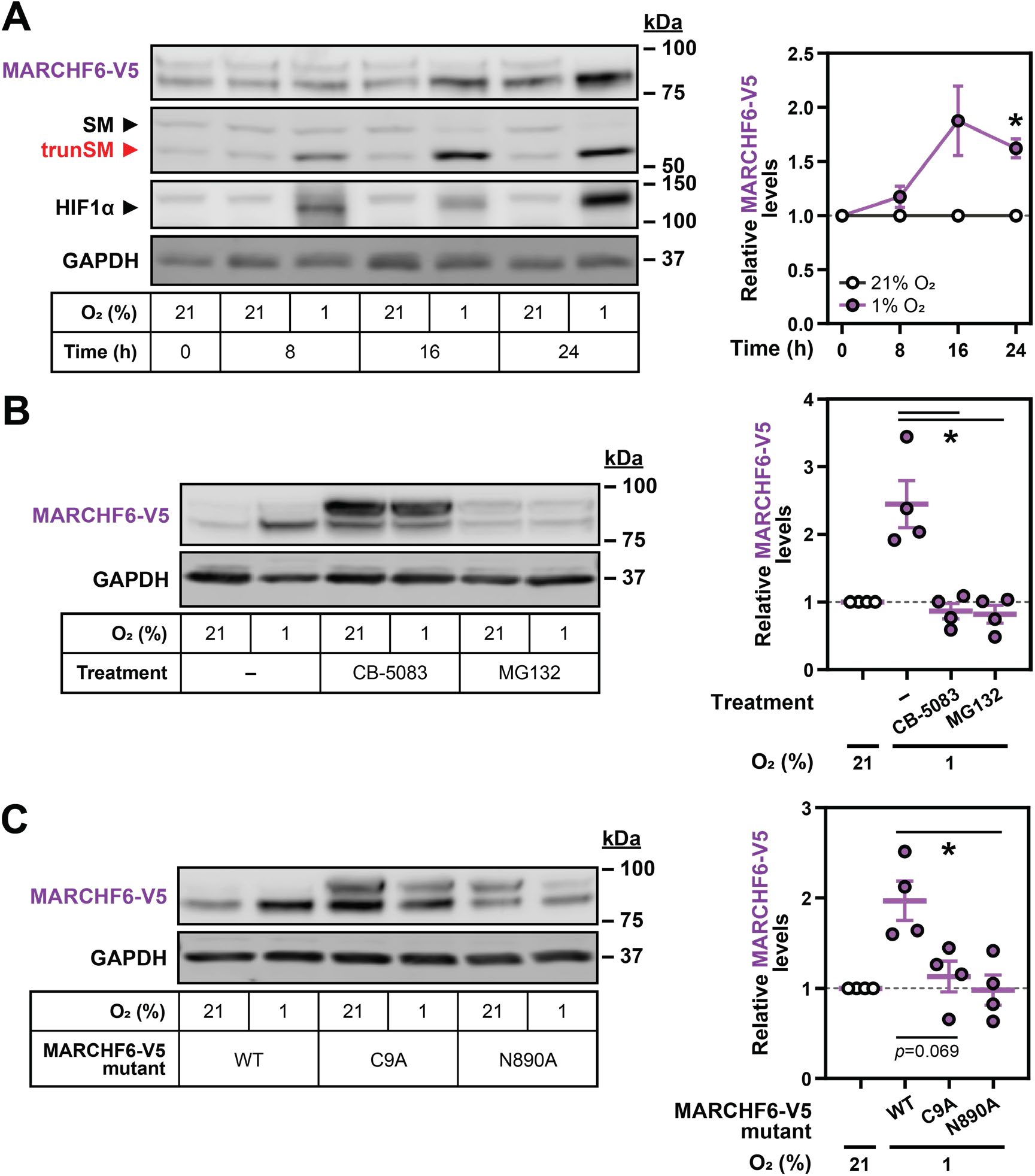
Hypoxia stabilizes the E3 ubiquitin ligase MARCHF6. **(A)** HEK MARCHF6-V5 cells were incubated under normoxic or hypoxic conditions for the indicated times. MARCHF6-V5 appears as two bands that were quantified collectively, as we have done previously [39]. **(B)** HEK MARCHF6-V5 cells were treated with or without 5 μM CB-5083 or 20 μM MG132 under normoxic or hypoxic conditions for 16 h. **(C)** The indicated HEK MARCHF6-V5 cell lines were incubated under normoxic or hypoxic conditions for 16 h. **(A–C)** Graphs depict densitometric quantification of MARCHF6-V5 levels normalized to the respective normoxic conditions, which were set to 1 (dotted line). Data presented as mean ± SEM from *n* ≥ 3 independent experiments (*, *p* ≤ 0.05; [A] two- tailed one-sample *t*-test vs. hypothetical mean of 1; [B, C] two-tailed ratio paired *t*-test vs. vehicle or WT condition).

Given that Asn-890 is required for the accumulation of MARCHF6 during hypoxia, we next considered whether this residue might be modified by asparagine hydroxylation. This is an oxygen-dependent modification known to control processes such as HIF1α target gene transactivation [42] and protein ubiquitination [43]. To determine if a similar mechanism regulates MARCHF6 in hypoxia, we used siRNA to deplete the asparaginyl hydroxylase FIH-1. Although this increased basal MARCHF6-V5 levels and blunted its hypoxia-induced accumulation (Supplementary Fig. S3A, B), confirmatory experiments using the FIH-1 inhibitor DM-NOFD [44] did not reproduce the effect (Supplementary Fig. S3C). We concluded that asparagine hydroxylation is unrelated to the reduced autoubiquitination of MARCHF6 in hypoxia, with the true mechanism currently unknown.

### Hypoxia-induced squalene accumulation promotes partial degradation of SM

Having established that SM undergoes increased proteasomal degradation during hypoxia, we next investigated how low oxygen levels favor its partial rather than complete proteolysis to produce trunSM. As there is extensive precedent for metabolic regulation of cholesterol synthesis enzymes and the pathway contains multiple oxygen-dependent reactions, we considered whether the accumulation of a pathway intermediate might be responsible for this phenomenon. Hypoxia-induced accumulation of trunSM occurred in cells incubated under both lipoprotein-replete and lipoprotein-deficient conditions, in which the cholesterol synthesis pathway is active (Fig. 4A). However, its accumulation was abolished when lipoprotein-deficient cells were co-treated with a statin to inhibit HMGCR and the early cholesterol synthesis pathway. By contrast, there was no effect of sterol depletion on the disappearance of full-length SM. This indicated that an intermediate or end-product of cholesterol synthesis promotes the partial rather than complete degradation of SM during hypoxia. We therefore turned our attention to the SM substrate squalene, given that it allosterically regulates SM degradation [8] and its conversion to monooxidosqualene is the first oxygen-dependent reaction of cholesterol synthesis. Squalene accumulated over the course of a hypoxic incubation (Fig. 4B, Supplementary Fig. S4), consistent with reduced SM activity under low-oxygen conditions. Moreover, this accumulation of squalene was strikingly well-correlated with the previously observed increase in trunSM levels (Fig. 4C), suggesting that the two responses may be linked.

**Figure 4.**
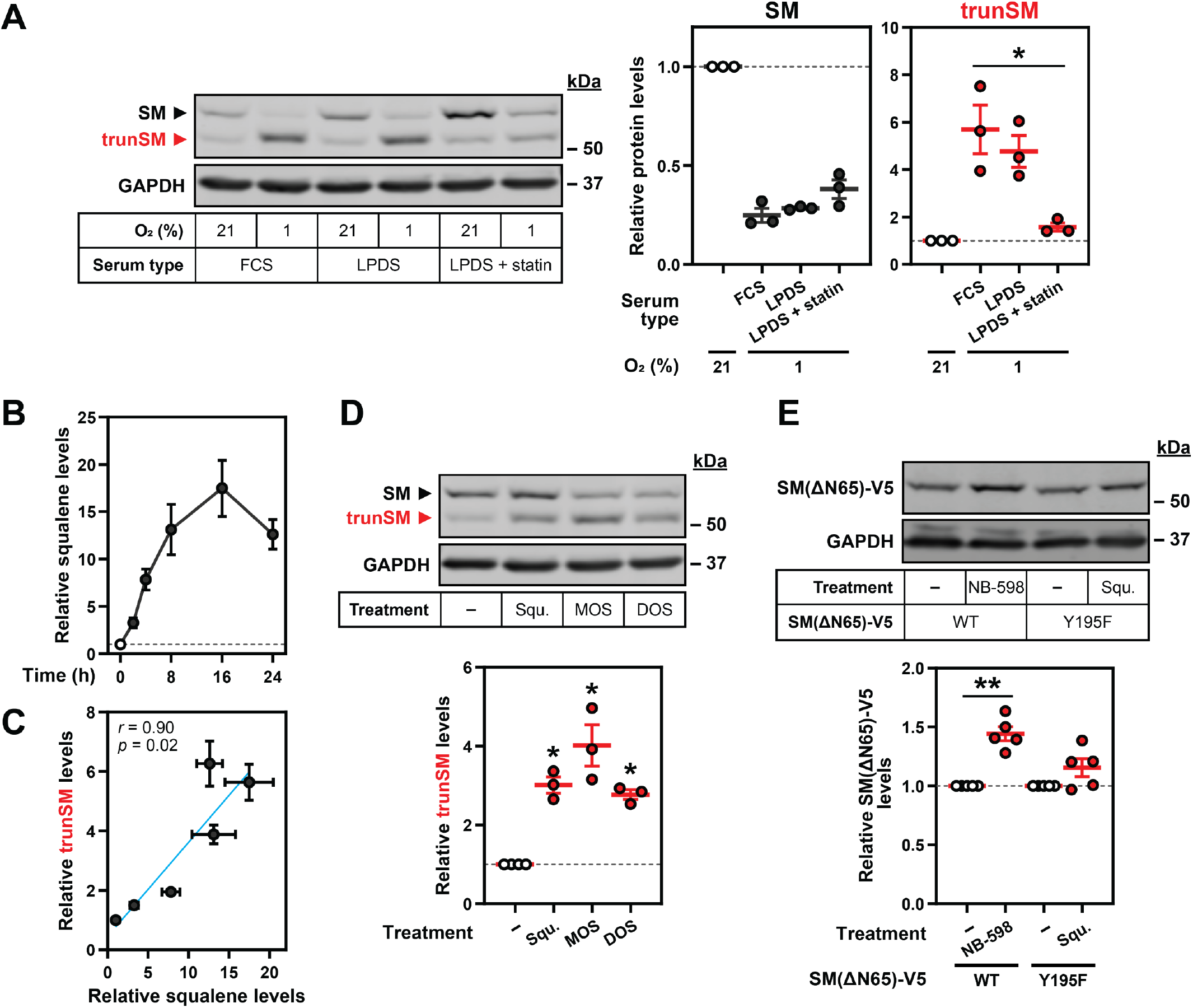
Hypoxia-induced squalene accumulation promotes partial degradation of SM. **(A)** HEK293T cells were incubated in medium containing fetal calf serum (FCS), lipoprotein-deficient FCS (LPDS) or LPDS containing 5 μM mevastatin and 50 μM mevalonolactone (LPDS + statin) for 8 h, refreshed in their respective media and incubated under normoxic or hypoxic conditions for 16 h. **(B)** HEK293T cells were incubated under normoxic or hypoxic conditions for the indicated times. Non-saponifiable lipids were extracted, and squalene levels were determined using gas chromatography-mass spectrometry and adjusted relative to the normoxic condition, which was set to 1 (dotted line). The maximal squalene level detected was 0.66 ± 0.12 ng per μg of total protein. **(C)** Pearson correlation between squalene levels in (B) and trunSM levels in Fig. 1C. Blue line indicates linear regression. **(D)** HEK293T cells were treated with or without 300 μM squalene (Squ.), monooxidosqualene (MOS) or dioxidosqualene (DOS) for 16 h. **(E)** HEK293T *SQLE*-knockout (*SQLE*-KO) clone 10 (c10) cells were transfected with the indicated constructs for 24 h, then treated with or without 1 μM NB-598 or 300 μM squalene for 16 h. **(A, D, E)** Graphs depict densitometric quantification of trunSM or truncated protein levels normalized to the (A) respective normoxic conditions or (D, E) respective vehicle conditions, which were set to 1 (dotted line). **(A–E)** Data presented as mean ± SEM from *n* ≥ 3 independent experiments (*, *p* ≤ 0.05; **, *p* ≤ 0.01; [A] two-tailed ratio paired *t*-test vs. FCS condition; [D, E] two-tailed one-sample *t*-test vs. hypothetical mean of 1).

Delivery of exogenous squalene mimicked the hypoxia-induced accumulation of trunSM in normoxic HEK293T (Fig. 4D, Supplementary Fig. S5A) and Huh7 cells (Supplementary Fig. S5B), confirming its ability to promote partial degradation of SM. Accumulation of trunSM was also induced by the oxygenated squalene derivatives monooxidosqualene and dioxidosqualene (Fig. 4D) but not by its saturated analogue squalane (Supplementary Fig. S5C), which has similar biophysical properties [45]. This indicated that truncation is promoted by squalene and its structurally related molecules in a specific manner, rather than through bulk membrane effects caused by lipid accumulation. To address the possibility that exogenous squalene is converted to a downstream product responsible for truncation, we performed squalene treatment of *SQLE*-knockout HEK293T cells (Supplementary Fig. S6A–C) transfected with a catalytically inactive SM Y195F mutant [46] to prevent the metabolism of added squalene. The truncated form of the Y195F mutant accumulated upon squalene treatment in *SQLE*-knockout cells, confirming that squalene alone can directly induce truncation (Supplementary Fig. S5D). There was no significant accumulation of the truncated fragment in cells transfected with wild-type SM, likely due to the clearance of exogenous squalene by the overexpressed protein and downstream enzymes.

To confirm that endogenously synthesized squalene is sufficient to trigger SM truncation, cells were treated with inhibitors of the relevant cholesterol synthesis enzymes (Supplementary Fig. S5A). The SM inhibitor NB-598 was excluded because of its ability to induce truncation through direct binding and stabilization of the SM catalytic domain that renders it resistant to proteasomal unfolding [8, 47]. Inhibition of squalene synthesis from farnesyl diphosphate (TAK-475) abolished the hypoxia-induced accumulation of trunSM, whereas elevated protein levels were still observed under conditions where squalene remained able to accumulate: inhibition of lanosterol synthesis from monooxidosqualene (BIBB 515), or inhibition of lanosterol demethylation (GR70585X) (Supplementary Fig. S5E). This result confirmed that lanosterol, which also accumulates during hypoxia [27], has no effect on SM truncation. We further noted that the inhibition of squalene or lanosterol synthesis, but not lanosterol demethylation, increased the levels of full-length SM and SM-N100-GFP-V5 under basal conditions. This was in line with our previous finding that farnesyl-containing molecules such as monooxidosqualene, dioxidosqualene and a squalene- derived probe can each stabilize SM *via* its regulatory domain in a similar manner to squalene itself [8]. Interestingly, the basal levels of trunSM were also increased in the presence of these inhibitors (Supplementary Fig. S5E), suggesting that all farnesyl-containing intermediates of cholesterol synthesis are capable of inducing SM truncation. Nevertheless, as the primary substrate of SM, squalene is likely to be the major driver of this process under hypoxic conditions.

SM contains two known squalene binding sites: the SM-N100 regulatory domain and the active site of the catalytic domain [8]. As direct binding of the SM inhibitor NB-598 to the catalytic domain results in truncation-promoting stabilization [9], we next considered whether squalene also exerts its effect on truncation through direct binding and stabilization of the catalytic domain. To eliminate the contribution of the SM-N100 domain, we transfected *SQLE*-knockout cells with an ectopic form of trunSM (SM[ΔN65]-V5) that was stabilized by NB-598 (Fig. 4E), consistent with past findings [8, 46]. By contrast, squalene treatment of cells expressing an inactive SM(ΔN65)-V5 mutant (Y195F) did not lead to significant protein stabilization. We concluded that squalene promotes SM truncation *via* the SM-N100 regulatory domain, rather than through direct binding and stabilization of the SM catalytic domain. Squalene treatment also did not increase MARCHF6-V5 levels (Supplementary Fig. S5F), suggesting that the stabilization of the E3 ubiquitin ligase during hypoxia is unrelated to squalene accumulation. This observation was consistent with the hypoxia-induced and MARCHF6-mediated degradation of full-length SM being unaffected by the activity of the cholesterol synthesis pathway (Fig. 4A). Together, this reinforced that squalene exerts its truncation-promoting effects post-ubiquitination, during the proteasomal- degradation of SM.

### SM truncation is increased in hypoxic endometrial cancer tissues

Aberrant SM expression and activity is an emerging hallmark of numerous malignancies [21, 48], yet the contribution of the constitutively active trunSM to oncogenesis is unknown. Therefore, we performed immunoblotting of paired tumor and adjacent benign tissue lysates from cohorts of lean and obese endometrial cancer patients. This cancer type was selected because of its close links with obesity [49, 50] and signaling by cholesterol-derived estrogens [51], as well as the unstudied status of SM in endometrial cancer. Similar trends were observed across both cohorts (Fig. 5A, Supplementary Fig. S7A–C), and the data were pooled. This showed that levels of HIF1α were increased in tumor tissues compared with adjacent benign tissues (Fig. 5B), consistent with the hypoxic conditions that occur within solid tumors [22]. Total SM levels were reduced in tumor tissues (Fig. 5B), yet this was accompanied by a dramatic increase in the proportion of SM that was truncated (Fig. 5C). Moreover, we observed a striking correlation between HIF1α levels and SM truncation across all tissues (Fig. 5D), further supporting the notion that hypoxia and truncation are linked. These results indicated that the hypoxia-induced truncation of SM is a physiologically relevant phenomenon in human tissues and may contribute to endometrial tumorigenesis.

**Figure 5.**
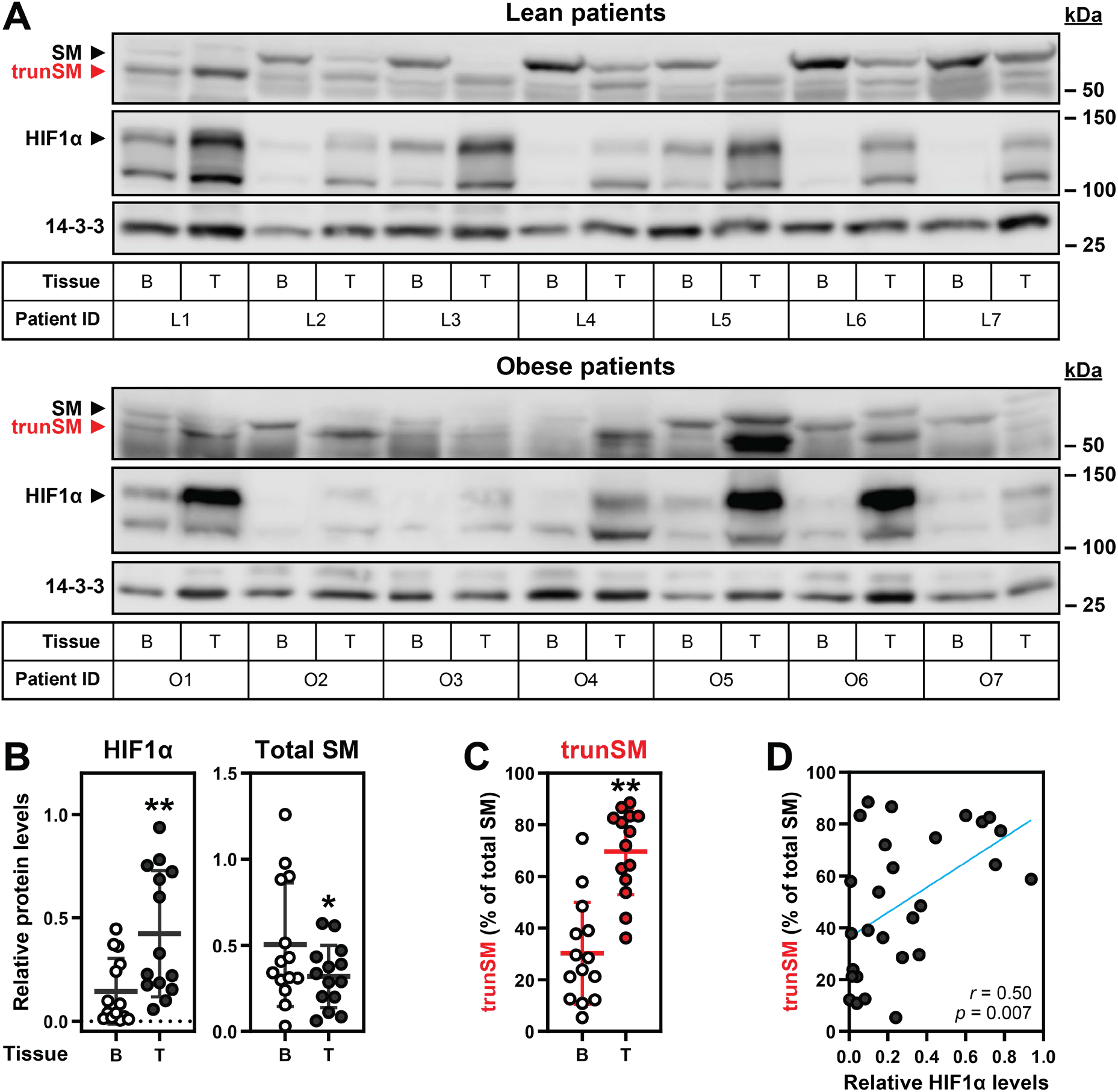
SM truncation is increased in hypoxic endometrial cancer tissues. **(A)** Paired tumor (T) and adjacent benign (B) tissue lysates from lean and obese endometrial cancer patients were analyzed by immunoblotting. **(B, C)** Graphs depict densitometric quantification of protein levels from (A) normalized to the 14-3-3 housekeeping protein, and (C) expressed as a proportion of total SM. Data presented as mean ± SD from *n* = 14 paired tissue sets (*, *p* ≤ 0.05; **, *p* ≤ 0.01; two-tailed paired *t*-test vs. adjacent benign tissue; normality of distributions confirmed by D’Agostino & Pearson testing [52]). **(D)** Pearson correlation between HIF1α levels in (B) and trunSM levels in (C). Blue line indicates linear regression.

## Discussion

Cholesterol synthesis is tightly regulated by metabolic supply and demand, with the lipid-sensing and rate-limiting enzyme SM being a key point at which this regulation is exerted. We previously showed that partial proteasomal degradation of SM produces a truncated and constructively active form of the enzyme [9]. In this study, we identified hypoxia as the physiological trigger for trunSM formation in both cell culture and human tissues *in situ* (Fig. 1). Hypoxia-induced truncation occurs through a two-part mechanism: (1) an increase in the levels of MARCHF6, the E3 ubiquitin ligase that targets SM to the proteasome (Fig. 2, Fig. 3), and (2) the accumulation of squalene, which impedes the complete degradation of SM and yields trunSM (Fig. 4). Beyond these mechanistic insights, we found that SM truncation is dramatically increased in endometrial cancer tissues and correlates with HIF1α levels (Fig. 5). Taken together, our results point towards the truncation of SM as an adaptive mechanism to promote squalene clearance during hypoxia, as well as a likely contributor to the widely reported oncogenic properties of SM.

### Cholesterol synthesis during hypoxia

Hypoxia places great strain on metabolic processes and necessitates the strict allotment of available oxygen and energy reserves. Cholesterol synthesis is a particularly resource-intensive pathway, requiring eleven oxygen molecules and over one hundred ATP equivalents per molecule of product, yet there are conflicting reports on changes in overall flux from acetyl-CoA to cholesterol during hypoxia [24, 25], indicative of cell type-specific responses. Nevertheless, the small number of studies into individual biosynthetic enzymes indicate that their activity is suppressed by hypoxia at multiple regulatory levels, which is supported by the accumulation of various pathway intermediates [26, 27, 53]. Lanosterol demethylase, which requires three oxygen molecules for catalysis, is transcriptionally downregulated by HIF2α and the hypoxia-induced long non-coding RNA *lincNORS*, contributing to the characteristic accumulation of lanosterol under hypoxic conditions [26, 54]. Lanosterol in turn triggers the ubiquitin-dependent degradation of the early cholesterol synthesis enzyme HMGCR [27], suppressing further oxygen consumption by the pathway. Our study expands this understanding of hypoxic adaptations by establishing that oxygen availability also regulates SM, a rate-limiting enzyme of cholesterol synthesis and the first to require molecular oxygen.

We found that SM is transcriptionally downregulated in hypoxic HEK293T cells but not MDA-MB-231 cells, consistent with previously reported cell-type specific changes in *SQLE* expression [23]. This accounted in part for the decline in full-length SM levels, although reduced *SQLE* translation through mechanisms such as mTOR suppression [55] cannot be ruled out as a contributing factor. We also found that, like HMGCR, the translated SM protein is targeted for proteasomal degradation during hypoxia. However, in stark contrast to HMGCR, the net effect of SM degradation under these conditions is the maintenance of, or even an increase in, the number of enzyme molecules available for catalysis. This is due to the increased partial proteolysis of SM to form trunSM, which lacks degron features within the SM-N100 regulatory domain and has a dramatically extended half-life yet remains catalytically active [9]. Disruption of the SM-N100 domain also renders trunSM resistant to cholesterol-induced degradation, which is an key element in the metabolic regulation of full-length SM [4, 9]. Therefore, the hypoxia-induced truncation of SM ensures that total enzyme levels remain constant even under cholesterol-replete conditions that would typically reduce its abundance.

The most plausible explanation for the seemingly paradoxical stabilization of SM during hypoxia is that this can compensate for reduced oxygen availability and allow enzyme activity to be maintained (Fig. 6). There are numerous ways in which the truncation and constitutive activity of SM is beneficial. During transient or low-level hypoxia, it may enable continued cholesterol synthesis so that cell growth and survival is not compromised by the oxygen shortfall. In support of this idea, hypoxia-induced cell death is exacerbated by SM inhibition [23]. Furthermore, the longevity of trunSM is likely to enable rapid resumption of pathway activity when normal oxygen levels are restored. During long-term or severe hypoxia, where there is insufficient oxygen for flux through downstream cholesterol synthesis, the role of trunSM in hypoxia may shift towards the efficient clearance of squalene. While generally considered an inert intermediate, excess squalene induces ER stress and is generally toxic in cells that lack SM activity or are unable to sequester squalene to lipid droplets [56, 57]. A secondary effect of squalene clearance is its downstream conversion to lanosterol, which promotes the shutdown of cholesterol synthesis *via* HMGCR degradation. This mechanism may explain why squalene accumulation lags behind that of lanosterol during the acute phase of hypoxia [27, 53]. Although we detected increased squalene levels at early hypoxic timepoints, a previous study of sterol-depleted cells did not [27], which is likely attributable to the basal upregulation of SM expression under these conditions. Elevated SM activity during hypoxia may also favor the synthesis of dioxidosqualene and ultimately 24(*S*),25-epoxycholesterol, a potent suppressor of cholesterol accretion [58]. However, this oxysterol is yet to be studied in a low-oxygen context.

**Figure 6.**
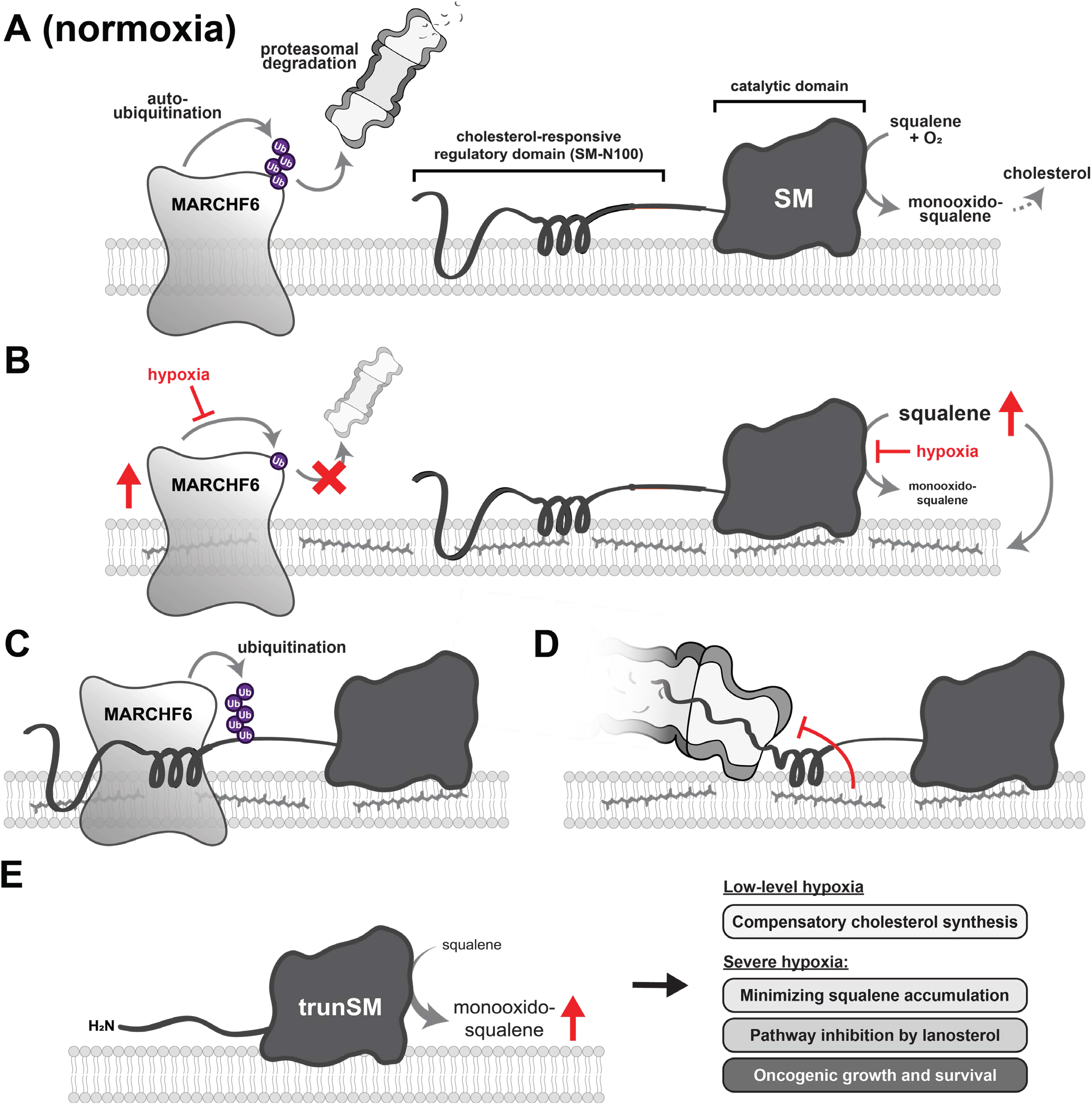
Model of hypoxia-induced SM truncation. **(A, B)** Hypoxic conditions inhibit the auto-ubiquitination of the E3 ubiquitin ligase MARCHF6, resulting in its stabilization. Simultaneously, the SM-catalyzed conversion of squalene to monooxidosqualene is inhibited, leading to squalene accumulation. **(C)** Increased MARCHF6 activity promotes the targeting of SM to the proteasome. **(D)** Squalene binds the SM-N100 domain and impedes the complete proteasomal degradation of SM, leading to **(E)** the formation of the constitutively active trunSM. During transient or low-level hypoxia, trunSM activity may facilitate continued cholesterol synthesis to compensate for the oxygen shortfall. During long-term or severe hypoxia, trunSM activity may reduce squalene-induced toxicity while also promoting the formation of lanosterol, which suppresses an early step of the cholesterol synthesis pathway. During pathophysiological hypoxia, trunSM may enable continued cell growth and survival *via* both cholesterol synthesis-dependent and -independent oncogenic properties.

### Molecular mechanism of truncation

The first step of SM truncation is its delivery to the proteasome [9]. This is enhanced during hypoxia by a novel ubiquitin signal at the Lys-82/90/100 cluster, as well as the accumulation of the E3 ubiquitin ligase MARCHF6, which facilitates the basal and metabolically-regulated degradation of SM [38]. Non-canonical ubiquitination sites required for cholesterol-induced degradation of SM [35] also contribute to its hypoxic degradation, which can be reconciled by the combination of increased MARCHF6 levels and the sterol-replete conditions under which these experiments were performed. Hypoxic accumulation of MARCHF6 occurs due to reduced auto-ubiquitination, as this effect is blocked by inhibition of the ER-associated degradation pathway or by mutation of residues essential for MARCHF6 activity. The precise mechanism by which low oxygen levels impede MARCHF6 auto-ubiquitination remains unclear, although the involvement of asparagine hydroxylation or accumulated squalene was ruled out. We previously found that cholesterol and several of its biosynthetic intermediates, including lanosterol, also impede MARCHF6 degradation [39]. This raises the possibility that a lanosterol-dependent mechanism promotes MARCHF6 accumulation during hypoxia, which is particularly intriguing given that MARCHF6 also targets lanosterol demethylase for degradation [59] and likely contributes to lanosterol accumulation. It remains to be determined how increased MARCHF6 levels influence the activity of its other metabolic substrates. These include the lipid droplet biogenesis protein perilipin-2 [60] and the thyroid hormone-activating deiodinase DIO2 [61], which both function in pathways perturbed by hypoxia [62, 63]. Unexpectedly, MARCHF6 levels were slightly but significantly reduced upon treatment with exogenous squalene. This was also observed for the saturated analogue squalane, and therefore is not a squalene-specific effect. Rather, it may suggest that the stability of the 14-transmembrane domain MARCHF6 protein is sensitive to membrane perturbations resulting from the delivery of large amounts of exogenous lipid.

A key finding of this study is that upon the hypoxia-induced delivery of SM to the proteasome, accumulated squalene inhibits its complete degradation and liberates the constitutively active trunSM. This feedforward mechanism, by which the substrate of SM preserves its activity, is mediated by the SM-N100 regulatory domain and functional under normoxic conditions. Here, it likely buffers against transient increases in squalene levels. Accumulation of trunSM occurs even when MARCHF6 is depleted, indicating that squalene similarly prevents the complete degradation of SM molecules targeted to the proteasome by other E3 ubiquitin ligases. These additional regulators, and the factors controlling their ubiquitination of SM, await future discovery and may shed light on how the SM-N100 regulatory domain undergoes MARCHF6-independent proteasomal degradation during hypoxia. Although hypoxia and lipid accumulation can trigger ER stress and altered proteostasis [64], the ability of squalene and farnesyl-containing compounds to allosterically bind the SM-N100 domain [8] strongly indicates that squalene induces SM truncation directly, rather than *via* bulk membrane effects. The specificity of this response is further supported by the inability of its saturated analog squalane to induce truncation. While squalene is the most likely of the farnesyl-containing cholesterol synthesis intermediates to accumulate under normal conditions, owing to the rate-limiting and oxygen-dependent nature of SM activity, our current and previous [8] data suggest that a farnesyl group alone is sufficient to promote truncation. Precisely how this lipophilic moiety interferes with SM degradation is unknown, although interaction with the membrane-embedded SM-N100 re- entrant loop [65] is a likely possibility. SM also contains a short intrinsically disordered domain (residues 81–120) that is required for its partial degradation and dictates the location of the truncation site [9], although interaction of a farnesyl group with this region is more difficult to envision.

Previously, we reported that squalene and farnesyl-containing compounds also blunt the MARCHF6-mediated ubiquitination of the SM-N100 domain, leading to stabilization of full-length SM [8]. The existence of dual mechanisms enabling squalene to sustain SM activity is intriguing, particularly as trunSM formation is itself dependent on ubiquitination [9]. One possibility is that truncation is stimulated at a lower squalene threshold than the inhibition of SM ubiquitination, allowing for a biphasic response to accumulating substrate. Another possibility is that truncation is a ‘failsafe’ mechanism to preserve the activity of the SM enzyme. This may be necessary when a reduced frequency of MARCHF6 ubiquitination is insufficient to completely prevent targeting of SM to the proteasome. It may also apply in situations where SM ubiquitination is promoted by other cellular stimuli. For instance, during hypoxia, the stabilization of MARCHF6 may override the inhibitory effects of accumulated squalene.

### Pathophysiological consequences of truncation

Overexpression and overactivity of SM occurs in a wide range of malignancies including hepatocellular carcinoma and prostate cancer, where it is positively correlated with severity and lethality [21, 48]. Despite this, a specific examination of trunSM levels had not been performed in any cancer type prior to our study. We found that total SM levels were reduced in endometrial tumor tissues, which was unexpected given that high *SQLE* mRNA expression is a risk factor for progression of this cancer [66]. However, the reduction in total SM was accompanied by a marked increase in the proportion of the stable and constitutively active trunSM. This was also well-correlated with the levels of HIF1α, suggesting that hypoxia-induced truncation of SM occurs *in vivo*. An increased proportion of trunSM was observed irrespective of whether patients were lean or obese, indicating that the phenomenon is likely not influenced by obesity or circulating lipid levels. Our data indicate that trunSM may be a contributing factor to cell growth and survival in the hypoxic tumor microenvironment, which is noteworthy given that severe hypoxia is often associated with a poorer cancer prognosis [22]. However, more work is needed to investigate the relationship between SM truncation and oncogenesis. There are currently no commercially available antibodies directed specifically against full-length SM, which precludes a comparison of its two forms using immunohistochemistry. Immunoblotting is thus the most robust means of determining trunSM levels. It would be worthwhile for future studies to quantify the truncation of SM in cancer types where its elevated protein expression has previously been identified, as well as in those that show a particular propensity for hypoxia, such as prostate and pancreatic cancer [28]. Although trunSM retains full catalytic activity and can likely fulfil the cholesterol synthesis-dependent roles of SM in oncogenesis, further study is required to confirm that it can enable the suite of other oncogenic functions previously reported for full-length SM. To date, these include the activation of ERK signaling [18] as well as interactions with carbonic anhydrase 3 [20], GSK3β and p53 [19]. These cholesterol-independent effects of SM are likely to be most important during severe hypoxia, where cholesterol synthesis cannot proceed.

Finally, although this study was largely performed using the extremely low oxygen levels characteristic of pathophysiological hypoxia, we also observed that SM truncation was responsive to a range of ‘physoxic’ oxygen levels that occur within normal tissues *in situ*. This reinforces the concept of truncation as a buffering mechanism to accommodate normal substrate fluctuations, which may be important for preventing squalene-induced dysfunction in tissues with lower oxygen perfusion, such as the brain [28]. Squalene fluctuations also occur diurnally [67] and with changing sterol status [4], and its levels can vary dramatically between different tissues [68]. Thus, the ability of squalene to stimulate formation of the rate-617 limiting trunSM may play an important role in regulating flux through cholesterol synthesis 618 under varying physiological conditions.

## Materials and methods

### Reagents and cell lines

Fetal calf serum (FCS), newborn calf serum (NCS), high-glucose Dulbecco’s Modified Eagle’s Medium (DMEM-HG), penicillin/streptomycin, Opti-MEM reduced serum medium, RNAiMAX transfection reagent, Lipofectamine 3000 transfection reagent, TRI reagent, the SuperScript III First-Strand Synthesis kit, and the PureLink Genomic DNA Mini kit were from Thermo Fisher (Carlsbad, CA, USA). Lipoprotein-deficient serum (LPDS, 30 mg/mL protein) was prepared from FCS by density gradient centrifugation and dialysis, as described in [69]. Primers, small interfering RNA (siRNA), protease inhibitor cocktail, and Tween-20 were from Sigma-Aldrich (St Louis, MO, USA). Tris-glycine SDS-PAGE gels were prepared in-house. Immobilon Western chemiluminescent HRP substrate and nitrocellulose membranes were from Millipore (Burlington, MA, USA). The QuantiNova SYBR Green PCR kit was from Qiagen (Hilden, GE). Skim milk powder was from Fonterra (Richmond, VIC, AU). Chemicals were from the following suppliers: 2,3,22,23-dioxidosqualene (Echelon Biosciences S-0302), 2,3-oxidosqualene (monooxidosqualene; Echelon Biosciences S-0301), 5α-cholestane (Sigma-Aldrich C8003), bafilomycin A1 (Sigma-Aldrich B1793), BIBB 515 (Cayman Chemical 10010517), CB-5083 (Cayman Chemical 16276), DM-NOFD (Sigma-Aldrich SML1874), DMOG (Sigma-Aldrich D3695), FG-4592 (Cayman Chemical 15294), GR70585X (GlaxoSmithKlein), mevalonolactone (Sigma-Aldrich M4667), mevastatin (Sigma-Aldrich M2537), MG132 (Sigma-Aldrich C2211), NB-598 (Chemscene CS-1274), squalane (Sigma-Aldrich 234311), squalene (Sigma-Aldrich S3626), TAK-475 (Sigma-Aldrich SML2168).

Human-derived HEK293T cells were a gift from the UNSW School of Medical Sciences (UNSW Sydney, NSW, Australia), HCT116 cells were a gift from Dr Ewa Goldys (UNSW Sydney, NSW, Australia), Huh7 cells were a gift from the Centre for Cardiovascular Research (UNSW Sydney NSW, Australia), and MBA-MB-231 and HeLaT cells were gifts from Dr Louise Lutze-Mann and Dr Noel Whitaker (UNSW Sydney, NSW, Australia). HEK293 cells stably expressing SM-N100-GFP-V5, MARCHF6-V5, MARCHF6-V5 C9A, or MARCHF6-V5 N890A were generated previously using the Flp-In T-Rex system [38, 39].

### Cell culture

Cells were maintained in a humidified Heraeus BB 15 incubator at 37 °C, 5% CO_2_, and 21% O_2_(normoxia) in maintenance medium (DMEM-HG, 10% [v/v] FCS, 100 U/mL penicillin, 100 μg/mL streptomycin). To improve HEK293 and HEK293T cell surface adhesion, culture vessels were treated with 25 μg/mL polyethyleneimine in phosphate- buffered saline (PBS) for 15 min at 37 °C prior to cell seeding. Plasmid and siRNA transfections were performed in maintenance medium lacking penicillin and streptomycin. Sterol depletions were performed in maintenance medium containing LPDS. Hypoxic incubations at 0.5–10% O_2_were performed in a humidified Binder CB 150 incubator at 37 °C and 5% CO_2_. For all treatments, appropriate solvent controls were used (water [DMOG]; ethanol (mevalonolactone, GR70585X); dimethyl sulfoxide [CB-5083, MG132, bafilomycin A1, DM-NOFD, mevastatin, TAK-475, BIBB-515, NB-598]; dimethyl sulfoxide containing 1% [v/v] Tween-20 [squalene, monooxidosqualene, dioxidosqualene, squalane]) and the final concentration of solvent did not exceed 1% (v/v) in cell culture medium. Treatments were delivered in full medium refreshes, and all experiments were 48–72 h in duration.

### Plasmids

Plasmids encoding Cas9 and *SQLE*-targeting guide RNAs were generated by *BbsI* cloning into a PX458 vector as described in [70]. Amino acid substitutions (SM K290R, Y195F) within expression vectors were generated using the overlap extension cloning method, as described previously [71]. The identity of all plasmids was confirmed *via* Sanger dideoxy sequencing. The plasmids used in this study are listed in Supplementary Table 1, and the primer sequences used for DNA cloning are listed in Supplementary Table 2.

### siRNA and plasmid transfection

To downregulate gene expression or transiently overexpress proteins, cells were seeded into 12-well plates. The next day, cells were transfected with 15 pmol siRNA using RNAiMAX (Invitrogen; 15 pmol siRNA: 2 μL reagent) or 1 μg expression vector using Lipofectamine 3000 (Invitrogen; 1 μg DNA: 2 μL reagent with 2 μL P3000 supplemental reagent), delivered in Opti-MEM. After 24 h, cells were refreshed in maintenance medium and treated as specified in figure legends. The siRNAs and plasmids used in this study are listed in Supplementary Table 1.

### Protein harvest and immunoblotting

To quantify protein levels, cells were seeded into 6- or 12-well plates and treated as specified in figure legends. For detection of SM, total protein was harvested in 2% SDS lysis buffer (10 mM Tris-HCl [pH 7.6], 2% [w/v] SDS, 100 mM NaCl, 2% [v/v] protease inhibitor cocktail), passed through a 21-gauge needle until homogenous, and vortexed at room temperature for 20 min. For detection of MARCHF6, cells were scraped in ice-cold PBS, pelleted, and lysed in modified RIPA buffer (50 mM Tris-HCl [pH 8.0], 0.1% [w/v] SDS, 1.5% [w/v] IGEPAL CA-630, 0.5% [w/v] sodium deoxycholate, 150 mM NaCl, 2 mM MgCl_2_, 2% [v/v] protease inhibitor cocktail), passed 20 times through a 22-gauge needle, rotated at 4 °C for 30 min, and centrifuged at 17,000 *g* and 4 °C for 15 min to obtain the supernatant. Lysate protein content was quantified using the bicinchoninic acid assay (Thermo Fisher), and sample concentrations were normalized by dilution in the appropriate lysis buffer and 0.25 vol 5× Laemmli buffer (250 mM Tris-HCl [pH 6.8], 10% [w/v] SDS, 25% [v/v] glycerol, 0.2% [w/v] bromophenol blue, 5% [v/v] β-mercaptoethanol). For SM detection, normalized samples were heated at 95°C for 5 min.

Proteins were separated by 10% (w/v) Tris-glycine SDS-PAGE, electroblotted onto nitrocellulose membranes, and blocked in 5% (w/v) skim milk powder in PBS containing 0.1% (v/v) Tween-20 (PBST). Immunoblotting was performed using the following antibodies: rabbit polyclonal anti-SM(SQLE) (Proteintech 12544-1-AP; 1:2,500 at 4 °C for 16 h), rabbit polyclonal anti-HIF1α (Proteintech 20960-1-AP; 1:1,000 at room temperature for 1 h), rabbit monoclonal anti-GAPDH (Cell Signaling Technology 2118; 1:2,000 at 4 °C for 16 h), mouse monoclonal anti-V5 (Thermo Fisher R960-25; 1:5,000 at room temperature for 1 h), mouse monoclonal anti-pan-14-3-3 (Santa Cruz Biotechnology sc-1657; 1:1,000 at 4 °C for 16 h), IRDye 680RD donkey anti-rabbit IgG (LI-COR Biosciences LCR-926-68073; 1:5,000), IRDye 800CW donkey anti-mouse IgG (LI-COR Biosciences LCR-926-32212; 1:10,000 at room temperature for 1 h), peroxidase-conjugated AffiniPure donkey anti-rabbit IgG (Jackson ImmunoResearch Laboratories 711-035-152; 1:10,000 at room temperature for 1 h), and peroxidase-conjugated AffiniPure donkey anti-mouse IgG (Jackson ImmunoResearch Laboratories 715-035-150; 1:10,000 at room temperature for 1 h). All antibodies were diluted in 5% (w/v) bovine serum albumin in PBST containing 0.02% (w/v) sodium azide, except for anti-HIF1α and peroxidase-conjugated antibodies, which were diluted in 5% (w/v) skim milk powder in PBST. Fluorescence-based detection of SM and trunSM, GAPDH, SM-N100-GFP-V5, and (HA)_3_-SM-V5 was performed using an Odyssey CLx imager (LI-COR Biosciences), and enhanced chemiluminescence-based detection of HIF1α, MARCHF6-V5, SM and trunSM (tissue lysates) and 14-3-3 was performed using Immobilon Western chemiluminescent HRP substrate (Millipore) and an ImageQuant LAS 500 imager (Cytiva Life Sciences). Densitometry analysis was performed using Image Studio Lite v5.2.5 (LI-COR Biosciences).

### RNA harvest and qRT-PCR

To quantify gene expression, cells were seeded into 12-well plates and treated as specified in figure legends. Total RNA was harvested using TRI reagent (Sigma-Aldrich) and polyadenylated RNA was reverse transcribed using the SuperScript III First Strand Synthesis kit (Invitrogen). cDNA products were used as the template for quantitative reverse transcription-PCR (qRT-PCR) in technical triplicate using the QuantiNova SYBR Green PCR kit (Qiagen). mRNA levels were normalized to the geometric mean of *RPL11*, *GAPDH* and *ACTB* for hypoxia experiments, or *PBGD* for validation of siRNA knockdowns and gene knockout, using the comparative C_T_method [72]. Normalized data were adjusted relative to the control condition, as specified in figure legends. The primer sequences used for qRT-PCR in this study are listed in Supplementary Table 2.

### Gas chromatography-mass spectrometry

To quantify cellular squalene levels, gas chromatography-mass spectrometry was performed as described previously [8]. Briefly, cells in 6-well plates were lysed in 0.05 M NaOH, total protein was quantified using the bicinchoninic acid assay, and samples were adjusted to the lowest protein concentration using 0.05 M NaOH plus 4 μg 5α-cholestane as an internal standard, in a total volume of 1 mL. Lysates were mixed with 1 mL 100% (v/v) ethanol, 500 μL 75% (w/v) KOH, 1 μL 20 mM butylated hydroxytoluene, and 20 μL 20 mM EDTA, and saponified at 70 °C for 1 h. Non-saponifiable lipids were extracted by mixing with 1 mL 100% (v/v) ethanol and 2.5 mL hexane, centrifuging at 4,000 *g* for 5 min, and collection of 2 mL of the organic phase. Lipids were dried in a vacuum centrifuge, resuspended in 50 μL *N*,*O*-bis(trimethylsilyl)trifluoroacetamide, and derivatized at 60 °C for 1 h.

Derivatized lipids (1.5 μL) were injected *via* a heated (300 °C) splitless with surge (38.0 psi for 0.50 min) inlet into a Thermo Trace gas chromatograph fitted with a Trace TR-50MS GC column (60 m × 0.25 mm, 0.25 μm film thickness, Thermo Fisher). Analytes were separated with helium as the carrier gas at a constant flow of 1.2 ml/min with vacuum compensation, and temperature programming as follows: 70 °C for 0.70 min, 20 °C/min to 250 °C, 3 °C/min to 270 °C, 1.5 °C/min to 315 °C, then hold for 10 min. The GC column was coupled to a Thermo DSQIII mass spectrometer, with a transfer line temperature of 320 °C and an ion source temperature of 250 °C. For mass spectrometry analysis, the electron energy was 70 eV, the emission current was 130 μA and the detector gain was 3.0 × 10^5^. Squalene and 5α-cholestane standards were analyzed in scan mode (34–600 Da) to identify peaks and retention times, and identity was confirmed using the National Institute of Standards and Technology databases. Experimental samples were analyzed in selective ion monitoring mode to detect squalene (*m*/*z* = 81.0, 410.4) and 5α-cholestane (*m*/*z* = 149.1, 217.2, 372.4), with a detection width of 0.1 and dwell time of 200 ms. Chromatograph peaks were integrated using Thermo Xcalibur software (v2.2 SP1.48) and the peak area of squalene was normalized to the 5α-cholestane internal standard. A squalene standard curve ranging from 3.125–100 ng/μL was used to quantify squalene levels, with data adjusted to the total protein content of the cell lysate.

### Generation of SM knockout cells

To knock out SM using CRISPR/Cas9, guide RNAs targeting the *SQLE* proximal promoter (hg38 chr8:124998048–124998067; AATGGAAACGTTCCGACCCG) and first exon (hg38 chr8:124999588–124999607; ATCCGAGAAGAGGGCGAACT) were designed using CHOPCHOP [73] and cloned into the Cas9- and GFP-encoding PX598 vector using BbsI restriction enzyme cloning, as described in [70]. HEK293T cells were seeded into 10 cm dishes and transfected with 5 μg of both vectors using Lipofectamine 3000, as described above. After 24 h, cells were trypsinized, washed with PBS and resuspended in FACS buffer (5% [v/v] FCS and 10 mM EDTA in PBS). Cells were sorted based on GFP fluorescence using a BD FACSAria III at the UNSW Flow Cytometry Facility. GFP-positive cells were seeded into 10 cm dishes at a low density (6,000–12,000 cells/dish) and allowed to adhere for 2–3 weeks. Single colonies were picked, expanded, and screened for *SQLE* mRNA expression and SM protein expression *via* qRT-PCR and immunoblotting, as described above. To confirm the genomic lesion, genomic DNA was isolated from clones using the PureLink Genomic DNA Mini kit (Thermo Fisher). The CRISPR/Cas9 target region was amplified and cloned into a pcDNA3.1 vector for Sanger dideoxy sequencing. The primer sequences used for guide RNA generation are listed in Supplementary Table 2.

### Endometrial cancer tissues

Matched tumor and adjacent benign endometrial tissues were obtained from lean (body mass index <27 kg/m^2^; *n* = 7) and obese (body mass index >30 kg/m^2^; *n* = 7) postmenopausal endometrioid adenocarcinoma patients recruited at the Royal Hospital for Women and Prince of Wales Private Hospital (Randwick, Sydney, Australia), as described in [74]. Consent was received from all patients prior to sample collection, and all processing and experiments were approved by the Human Research Ethics Committee (HREC) of the South Eastern Sydney Local Health District (HREC 15/339). Protein was isolated from powdered tissues using the AllPrep DNA/RNA/Protein Mini Kit (Qiagen). Protein pellets were lysed in HES-SDS buffer (20 mM HEPES [pH 7.4], 2% [w/v] SDS, 250 mM sucrose, 1 mM EDTA) and mixed with 1 vol 10% (w/v) SDS. Samples were homogenized using a handheld homogenizer and centrifuged at 15,000 *g* for 5 min. The protein content of the clarified supernatant was quantified using the bicinchoninic acid assay. Samples were diluted in 0.25 vol 5× Laemmli buffer and heated at 65 °C for 5 min prior to SDS-PAGE and immunoblotting.

### Data analysis and presentation

Data were normalized as described in figure legends. All data were obtained from *n* ≥ 3 independent experiments (biological replicates), and visualization and statistical testing were performed using GraphPad Prism v9.0 (GraphPad Software Inc.) as specified in figure legends. Where multiple statistical tests were conducted within a single experiment, *p*-values were corrected using the Benjamini-Hochberg method [75] with a false discovery threshold of 5%. Thresholds for statistical significance were defined as: *, *p* ≤ 0.05; **, *p* ≤ 0.01. Values of 0.05 < *p* < 0.075 are indicated in text on graphs. Figures were assembled using Adobe Illustrator v26.3 (Adobe Inc.).

## Acknowledgements

We thank Isabelle Capell-Hattam for assistance with CRISPR/Cas9, Dr Martin Bucknall for technical assistance with GC/MS, and the members of the Brown laboratory for critically reviewing this manuscript. We are also grateful to the patients from whom tissue samples were collected, and staff at the Royal Hospital for Women and Prince of Wales Private Hospital (Randwick, NSW, Australia) who assisted with surgical procedures. H.W.C. is supported by an Australian Research Training Program scholarship, R.F. is supported by a RANZCOG Mary Elizabeth Courier Research Scholarship, F.L.B. is supported by a Cancer Institute NSW Career Development Fellowship (2021/CDF1120), and the Brown laboratory is supported by Australian Research Council Grant DP170101178 and an NSW Health Investigator Development Grant.

## Author contributions

H.W.C.: conceptualization, methodology, investigation, visualization, writing – original draft. E.M.O.: investigation. X.D.: methodology, resources. R.F.: resources, funding acquisition. H.Y.: resources, funding acquisition. F.L.B.: resources, writing – review and editing, funding acquisition. A.J.B.: conceptualization, methodology, resources, writing – review and editing, supervision, funding acquisition.

---

**Supplementary Table S1.** siRNA and plasmids used for transfection.

| siRNA | Description |
| --- | --- |
| SIC001 | MISSION® universal negative control #1 |
| SASI_Hs01_00105239 | Targets human *MARCHF6* (NM_005885) |
| SASI_Hs01_00332063 | Targets human *HIF1A* (NM_001530) |
| SASI_Hs01_00019152 | Targets human *EPAS1 (HIF2A)* (NM_001430) |
| SASI_Hs01_00016727 | Targets human *HIF1AN (FIH-1)* (NM_017902) |
| Plasmid | Description |
| pCMV-(HA)<sub>3</sub>-SM-V5 | pcDNA3.1/V5-His TOPO vector containing the coding sequence of human squalene monooxygenase (SM; NM_003129.4) fused with three N-terminal HA epitope tags and C-terminal V5 and 6×His epitope tags, under the transcriptional control of a constitutive cytomegalovirus (CMV) promoter. Generated previously [9]. |
| pCMV-(HA)<sub>3</sub>-SM-V5 K82/90/100R | pCMV-(HA)<sub>3</sub>-SM-V5 containing SM K82R, K90R and K100R substitutions, which blunt SM truncation [9]. Generated previously [9]. |
| pCMV-(HA)<sub>3</sub>-SM-V5 K290R | pCMV-(HA)<sub>3</sub>-SM-V5 containing a K290R substitution, which disrupts a known ubiquitination site [34]. Generated during this study. |
| pCMV-(HA)<sub>3</sub>-SM-V5 SM-N100 C/S/T>A | pCMV-(HA)<sub>3</sub>-SM-V5 containing SM T3A, T9A, T11A, S43A, C46A, S59A, S61A, S67A, S71A, S83A and S87A substitutions, which blunt cholesterol-induced degradation of SM [35]. Generated previously [9]. |
| pCMV-(HA)<sub>3</sub>-SM-V5 Y195F | pCMV-(HA)<sub>3</sub>-SM-V5 containing an SM Y195F substitution that causes loss of catalytic activity [46]. Generated previously [9]. |
| pCMV-SM(ΔN65)-V5 | pCMV-(HA)<sub>3</sub>-SM-V5 containing a deletion of the N-terminal (HA)<sub>3</sub> tag and first 65 residues of SM, which are lost upon truncation [9]. Generated previously [9]. |
| pCMV-SM(ΔN65)-V5 Y195F | pCMV-SM(ΔN65)-V5 containing an SM Y195F substitution that causes loss of catalytic activity [46]. Generated during this study. |

**Supplementary Table S2.**
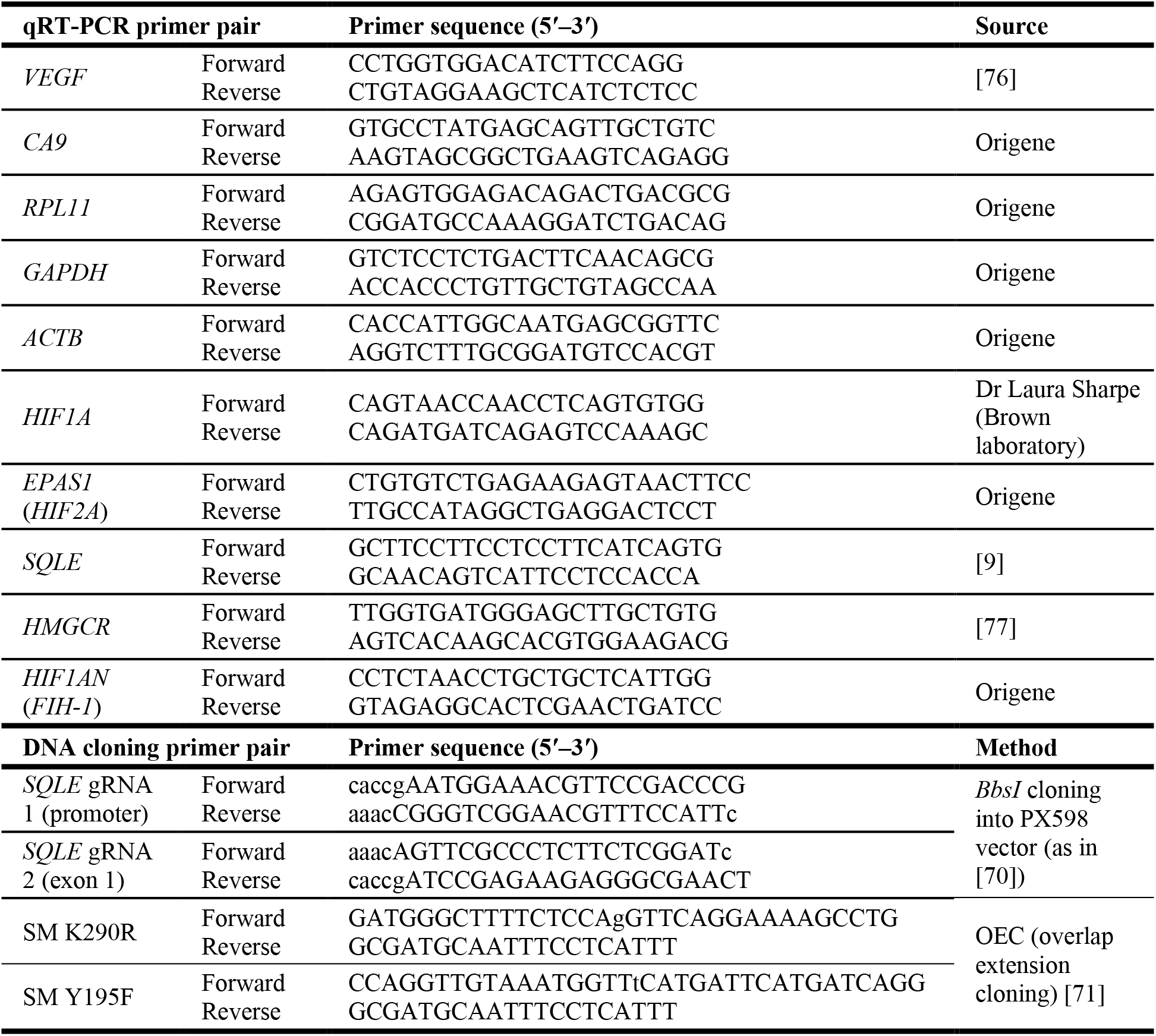
Primers used for qRT-PCR and DNA cloning. Non-annealing nucleotides for restriction enzyme cloning and DNA substitution are indicated in lowercase.

**Supplementary Figure S1.**
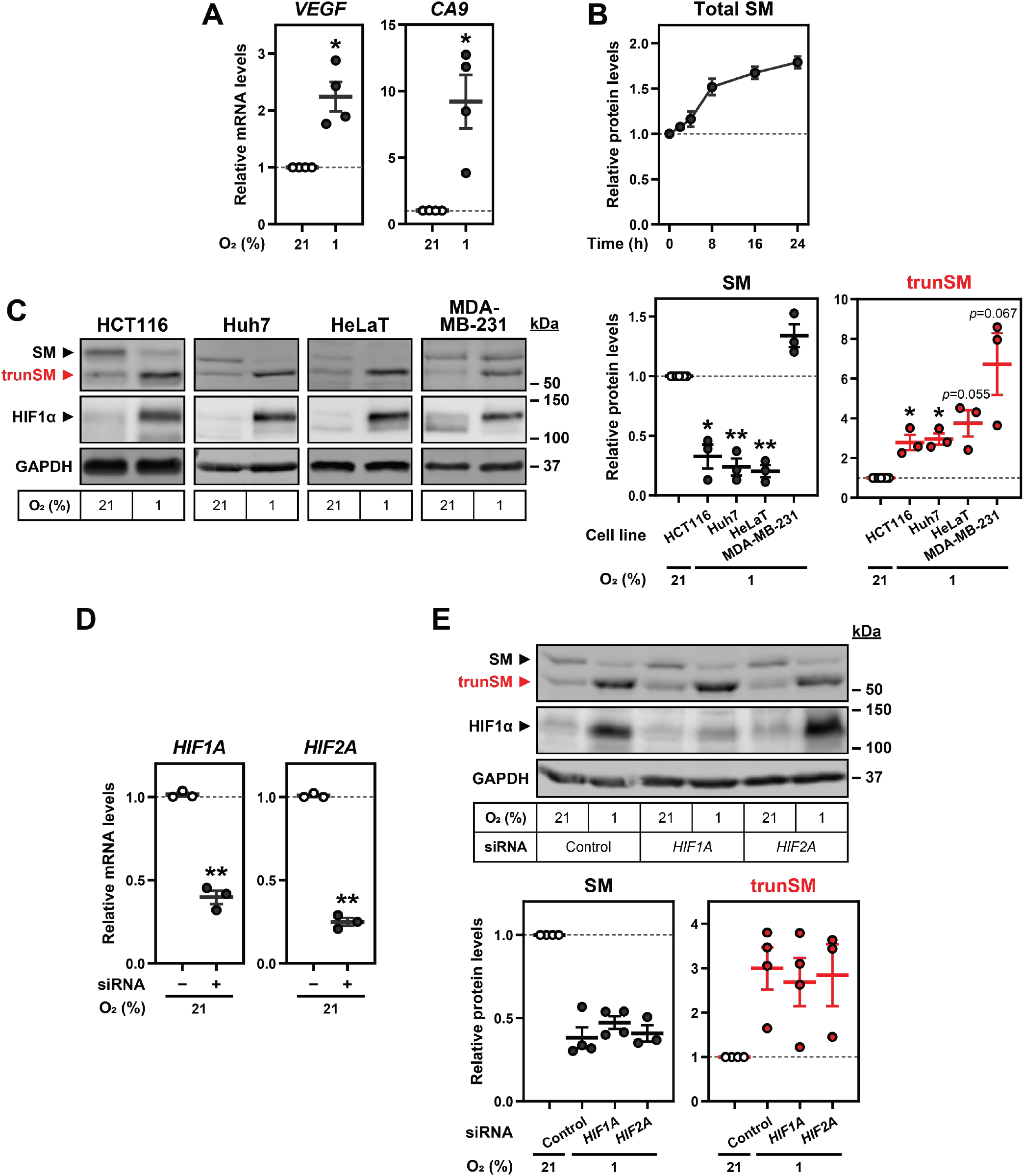
Hypoxia-induced truncation of SM is generalizable and independent of hypoxia-inducible factors.

**(A)** HEK293T cells were incubated under normoxic (21% O_2_) or hypoxic (1% O_2_) conditions for 24 h. Levels of the indicated HIF1α target gene transcripts were quantified, normalized to the levels of three housekeeping transcripts and adjusted relative to the normoxic condition, which was set to 1 (dotted line). **(B)** Quantification of total SM levels, expressed as the sum of SM and trunSM levels, in Fig. 1C. **(C)** The indicated cell lines were incubated under normoxic or hypoxic conditions for 24 h. **(D)** HEK293T cells were transfected with the indicated siRNAs for 24 h and incubated under normoxic or hypoxic conditions for 24 h. Levels of siRNA target transcripts in normoxic cells were quantified, normalized to the levels of the *PBGD* housekeeping transcript and adjusted relative to the control siRNA condition, which was set to 1 (dotted line). **(E)** HEK293T cells were treated as described in (B). **(B, C, E)** Graphs depict densitometric quantification of protein levels normalized to the respective normoxic conditions, which were set to 1 (dotted line). **(A–E)** Data presented as mean ± SEM from *n* ≥ 3 independent experiments (*, *p* ≤ 0.05; **, *p* ≤ 0.01; [A, C, D] two- tailed one-sample *t*-test vs. hypothetical mean of 1; [E] two-tailed ratio paired *t*-test vs. control siRNA condition).

**Supplementary Figure S2.**
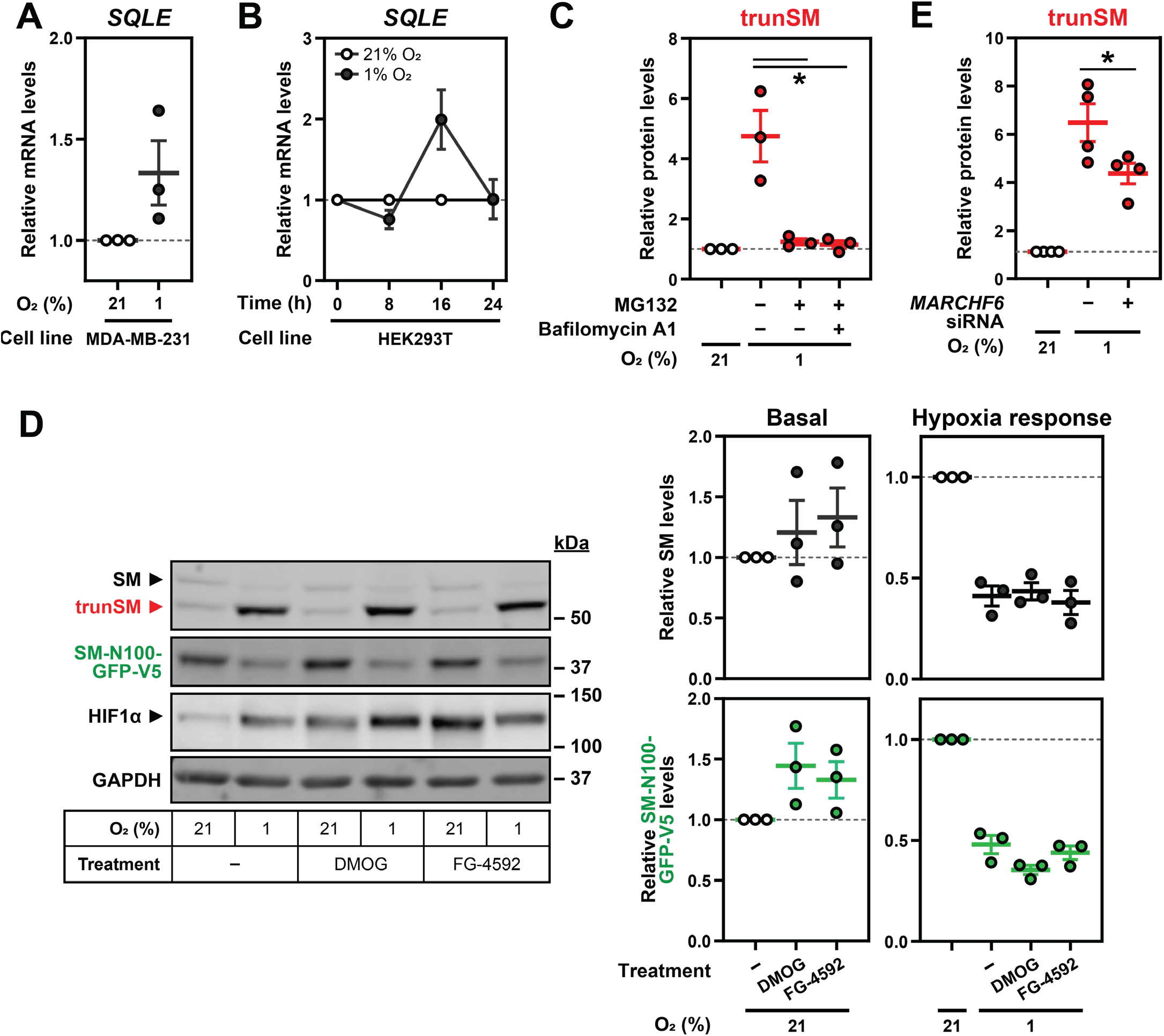
Hypoxia-induced degradation of full-length SM is independent of proline hydroxylation. **(A)** MDA-MB-321 cells were incubated under normoxic or hypoxic conditions for 24 h. Levels of *SQLE* transcripts were quantified, normalized to the levels of three housekeeping transcripts and adjusted relative to the normoxic condition, which was set to 1 (dotted line). **(B)** HEK293T cells were treated as described in Fig. 2B. Levels of *SQLE* transcripts were quantified, normalized to the levels of three housekeeping transcripts and adjusted relative to the normoxic condition, which was set to 1 (dotted line). **(C)** Quantification of trunSM levels in Fig. 2C. **(D)** HEK SM-N100-GFP-V5 cells were treated with or without 1 mM DMOG or 25 μM FG-4592 under normoxic or hypoxic conditions for 16 h. **(E)** Quantification of trunSM levels in Fig. 2E. **(C–E)** Graphs depict densitometric quantification of protein levels normalized to the respective normoxic conditions, which were set to 1 (dotted line). **(A–E)** Data presented as mean ± SEM from *n* ≥ 3 independent experiments (*, *p* ≤ 0.05; [A, B] two- tailed one-sample *t*-test vs. hypothetical mean of 1; [C–E] two-tailed ratio paired *t*-test vs. vehicle or control siRNA condition).

**Supplementary Figure S3.**
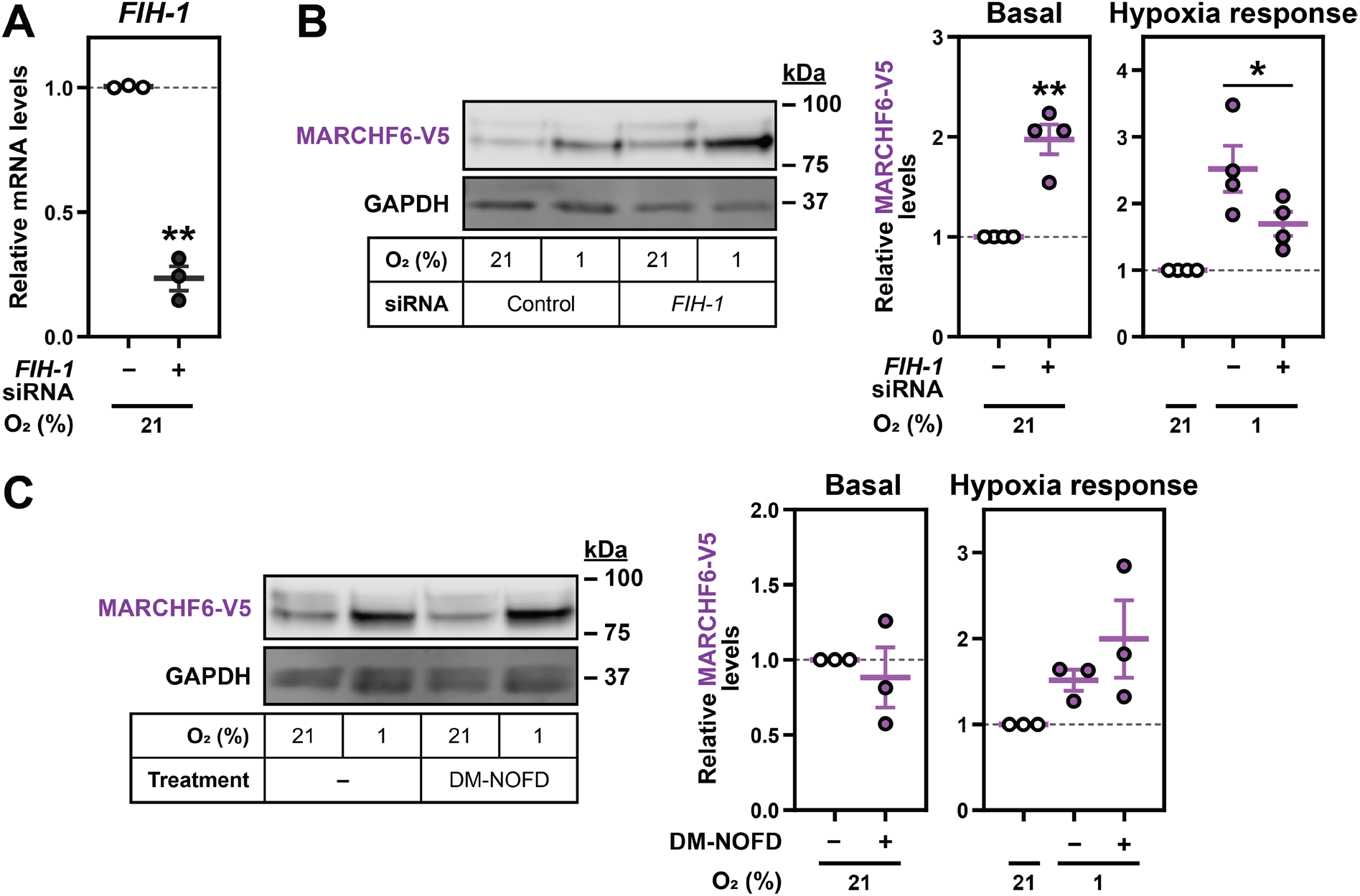
Hypoxia-induced stabilization of MARCHF6 is independent of asparagine hydroxylation. **(A)** HEK293T cells were transfected with control or *FIH-1* siRNA for 24 h and incubated under normoxic or hypoxic conditions for 24 h. Levels of *FIH-1* target transcripts in normoxic cells were quantified, normalized to the levels of the *PBGD* housekeeping transcript and adjusted relative to the control siRNA condition, which was set to 1 (dotted line). **(B)** HEK MARCHF6-V5 cells were transfected with control or *FIH-1* siRNA for 24 h and incubated under normoxic or hypoxic conditions for 16 h. Graphs depict densitometric quantification of MARCHF6-V5 levels normalized to the (left) vehicle or (right) normoxic condition, which was set to 1 (dotted line). **(C)** HEK MARCHF6-V5 cells were treated with 500 μM DM-NOFD under normoxic or hypoxic conditions for 16 h. **(B, C)** Graphs depict densitometric quantification of MARCHF6-V5 levels normalized to the respective normoxic conditions, which were set to 1 (dotted line). **(A–C)** Data presented as mean ± SEM from *n* ≥ 3 independent experiments (*, *p* ≤ 0.05; **, *p* ≤ 0.01; two-tailed one-sample *t*-test vs. hypothetical mean of 1 or [B, C] two-tailed ratio paired *t*-test vs. control siRNA or vehicle condition).

**Supplementary Figure S4.**
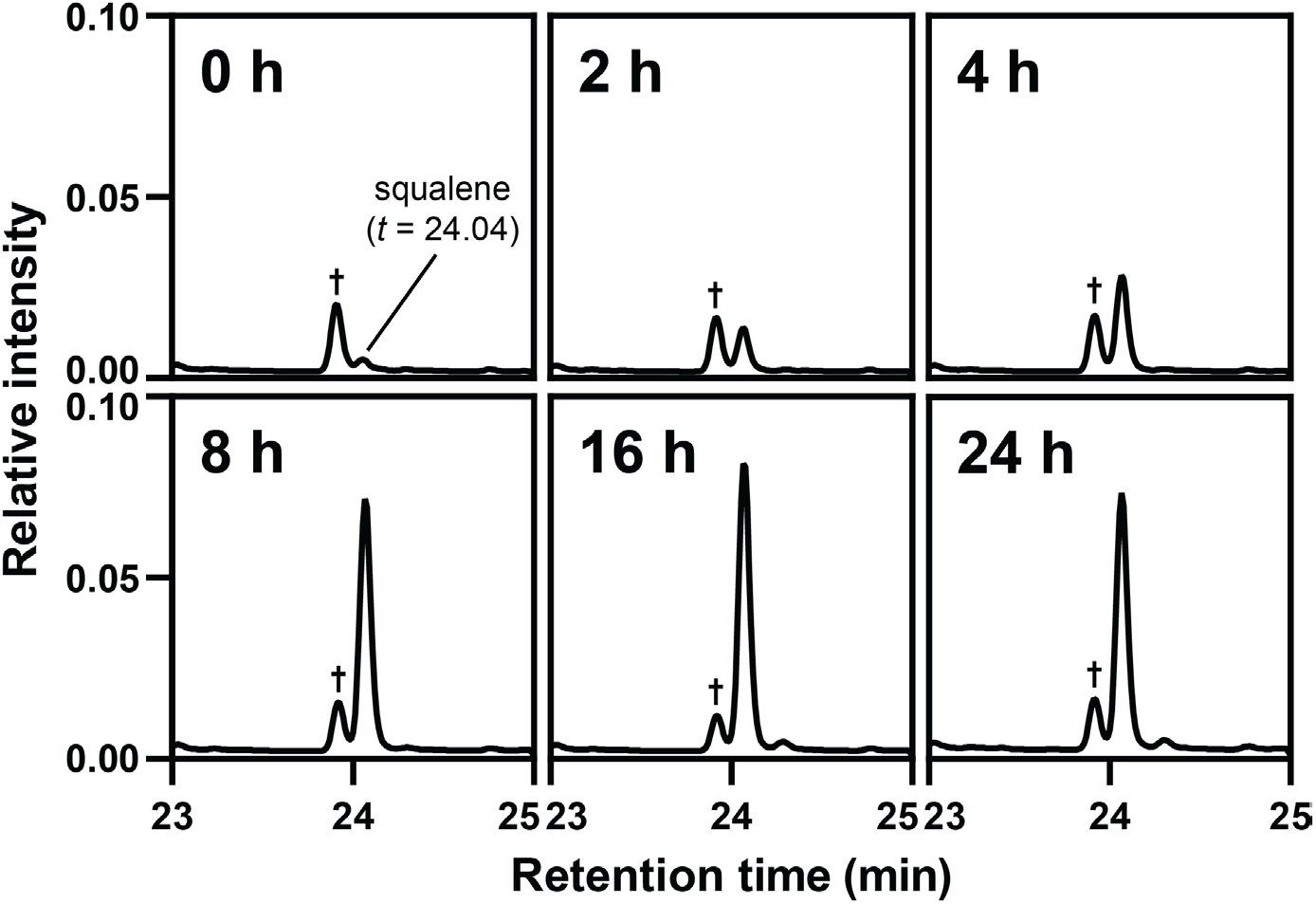
Hypoxia induces squalene accumulation. Representative selective ion monitoring chromatogram traces from gas chromatography-mass spectrometry analysis of squalene levels in Fig. 4B. Abundance was normalized to the 5α-cholestane internal standard, which was set to 1. Dagger indicates a non-specific analyte.

**Supplementary Figure S5.**
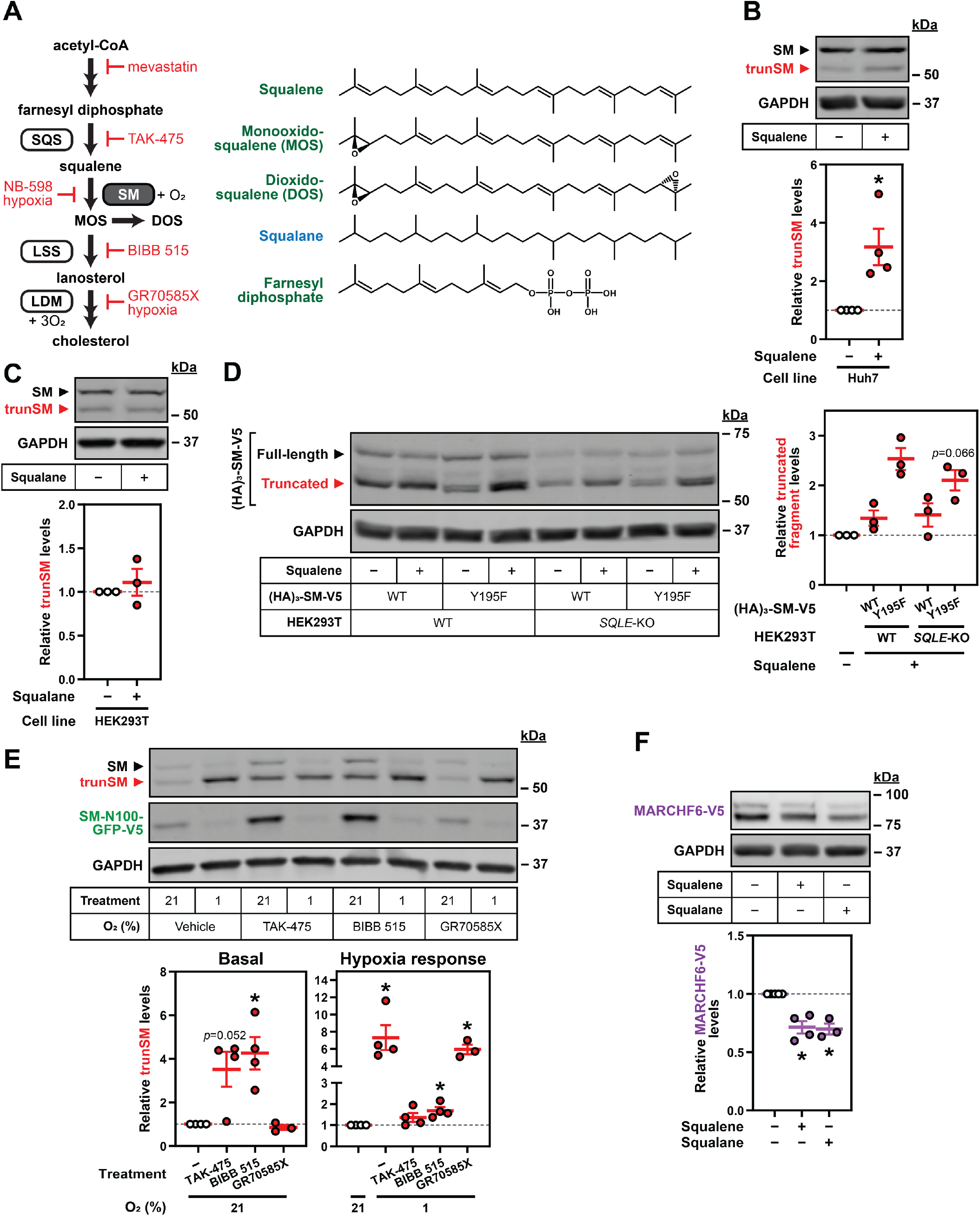
Farnesyl-containing cholesterol synthesis intermediates specifically promote partial degradation of SM. **(A)** Simplified schematic of the cholesterol synthesis pathway depicting activities of squalene synthase (SQS), SM, lanosterol synthase (LSS) and lanosterol 14α-demethylase (LDM), alongside chemical structures of squalene, monooxidosqualene (MOS), dioxidosqualene (DOS), squalane, and farnesyl diphosphate. Double-headed arrows indicate multiple enzymatic steps, red labels indicate inhibitors of cholesterol synthesis, and green and blue labels indicate molecules that were determined to promote or not to promote trunSM accumulation, respectively. **(B)** Huh7 cells were treated with or without 300 μM squalene for 16 h. **(C)** HEK293T cells were treated with or without 300 μM squalane for 16 h. **(D)** Parental (WT) or HEK293T *SQLE*-knockout (*SQLE*-KO) clone 10 (c10) cells were transfected with the indicated constructs for 24 h, then treated with or without 300 μM squalene for 16 h. **(E)** HEK SM-N100-GFP-V5 cells were treated with the indicated compounds under normoxic or hypoxic conditions for 16 h. **(F)** HEK MARCHF6-V5 cells were treated with or without 300 μM squalene or squalane for 16 h. **(B–F)** Graphs depict densitometric quantification of protein levels normalized to the respective vehicle or (E) respective normoxic conditions, which were set to 1 (dotted line). Data presented as mean ± SEM from *n* ≥ 3 independent experiments (*, *p* ≤ 0.05; two-tailed one-sample *t*-test vs. hypothetical mean of 1).

**Supplementary Figure S6.**
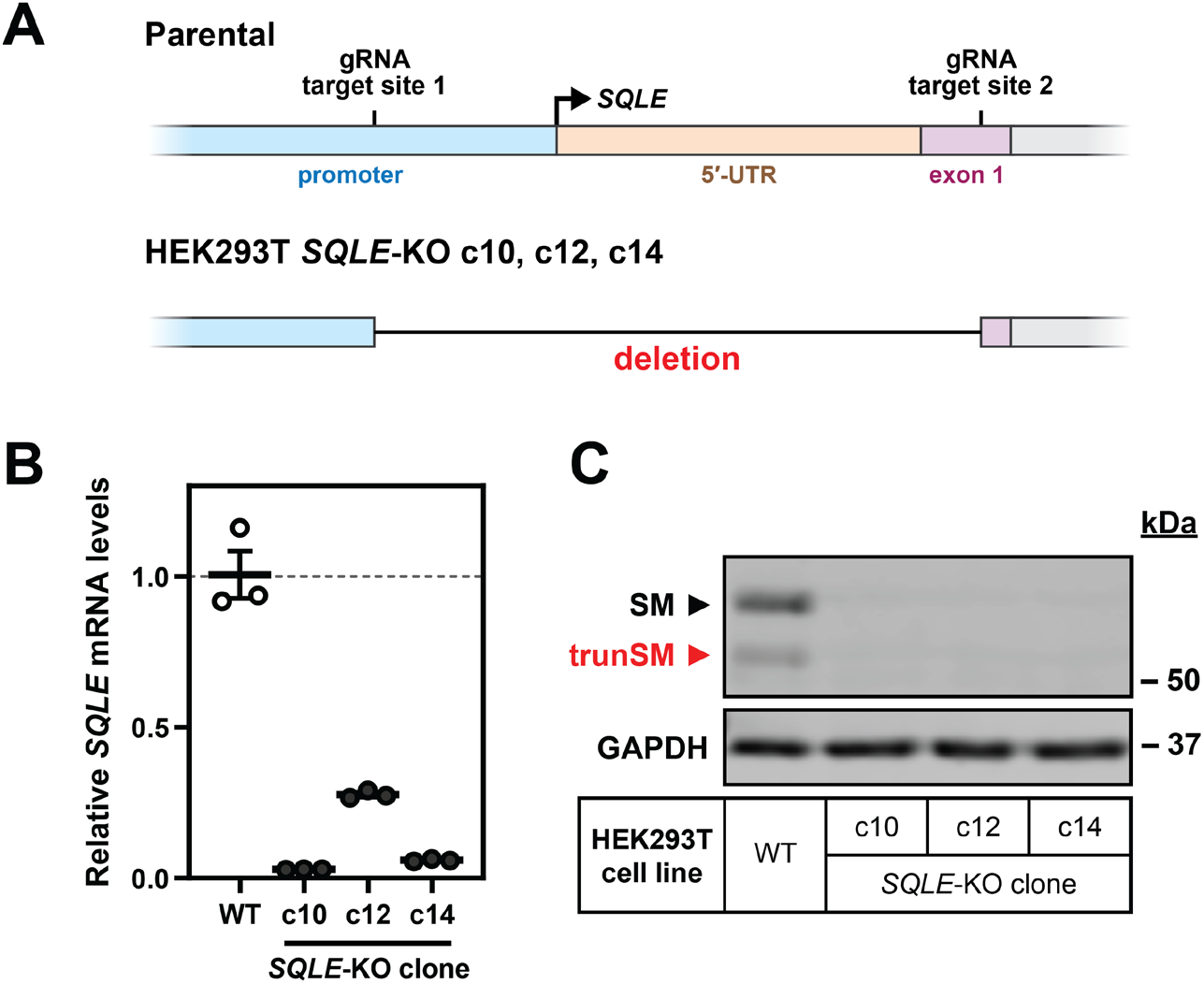
Generation of ***SQLE***-knockout HEK293T cells. **(A)** HEK293T cells were transfected with CRISPR/Cas9 guide RNAs (gRNAs) targeting the proximal promoter and first exon of the *SQLE* gene as indicated, and three clonal populations of *SQLE*-KO cells were isolated and expanded for each parental cell line. The target region was amplified from genomic DNA and sequenced, which found that all three clones contained a genomic deletion between the gRNA target sites. **(B)** *SQLE* transcript levels were quantified in HEK293T *SQLE*-KO clones, normalized to the *PBGD* housekeeping transcript, and adjusted relative to the parental (WT) cell line, which was set to 1 (dotted line). Data presented as mean ± SEM for technical triplicates from a single experiment. **(C)** Protein lysates from the HEK293T WT cell line and *SQLE*-KO clones were immunoblotted for SM and trunSM.

**Supplementary Figure S7.**
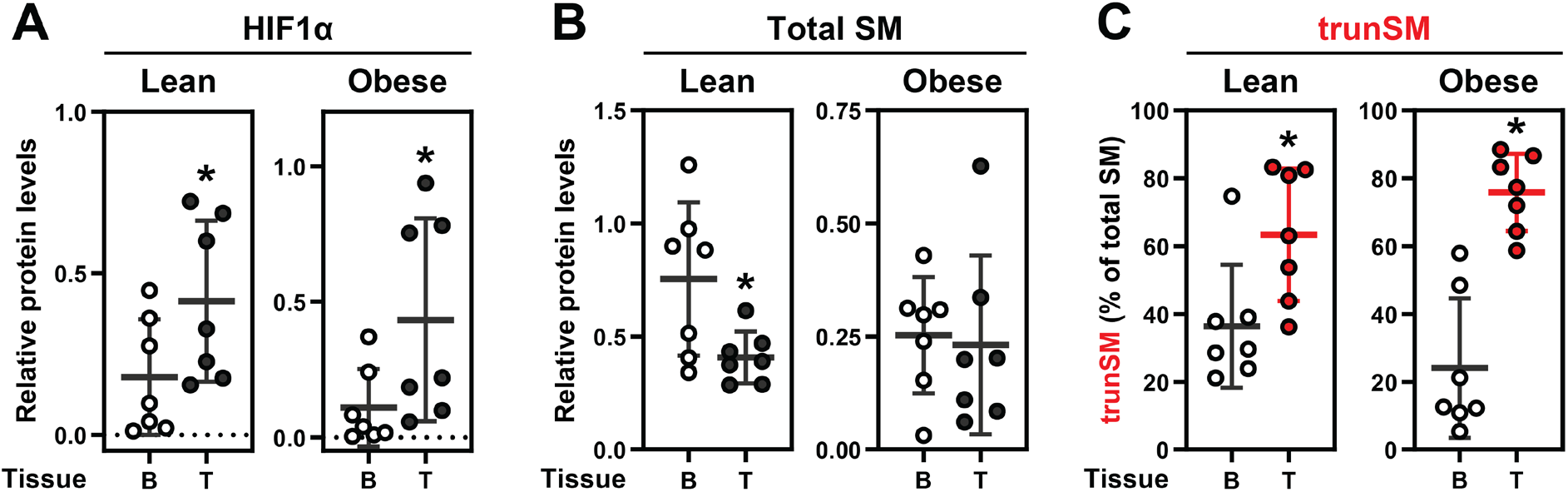
SM truncation is increased in endometrial cancer tissues from both lean and obese patients. Separate quantification of **(A)** HIF1α, **(B)** total SM, and **(C)** trunSM protein levels in tumor(T) and adjacent benign (B) tissues from lean and obese cohorts of endometrial cancer patients in Fig. 5A. Data presented as mean ± SD from *n* = 7 paired tissue sets in each cohort (*, *p* ≤ 0.05; two-tailed Wilcoxon matched-pairs signed rank test vs. adjacent benign tissue).

